# Neuraminidase inhibitors rewire neutrophil function *in vivo* in murine sepsis and *ex vivo* in COVID-19

**DOI:** 10.1101/2020.11.12.379115

**Authors:** Rodrigo de Oliveira Formiga, Flávia C. Amaral, Camila F. Souza, Daniel A. G. B. Mendes, Carlos W. S. Wanderley, Cristina B. Lorenzini, Adara A. Santos, Juliana Antônia, Lucas F. Faria, Caio C. Natale, Nicholas M. Paula, Priscila C. S. Silva, Fernanda R. Fonseca, Luan Aires, Nicoli Heck, Márick R. Starick, Celso M. Queiroz-Junior, Felipe R. S. Santos, Filipe R. O. de Souza, Vivian V. Costa, Shana P. C. Barroso, Alexandre Morrot, Johan Van Weyenbergh, Regina Sordi, Frederico Alisson-Silva, Fernando Q. Cunha, Edroaldo L. Rocha, Sylvie Chollet-Martin, Maria Margarita Hurtado-Nedelec, Clémence Martin, Pierre-Régis Burgel, Daniel S. Mansur, Rosemeri Maurici, Matthew S. Macauley, André Báfica, Véronique Witko-Sarsat, Fernando Spiller

## Abstract

Neutrophil overstimulation plays a crucial role in tissue damage during severe infections. Neuraminidase (NEU)-mediated cleavage of surface sialic acid has been demonstrated to regulate leukocyte responses. Here, we report that antiviral NEU inhibitors constrain host NEU activity, surface sialic acid release, ROS production, and NETs released by microbial-activated human neutrophils. *In vivo*, treatment with Oseltamivir results in infection control and host survival in peritonitis and pneumonia models of sepsis. Single-cell RNA sequencing re-analysis of publicly data sets of respiratory tract samples from critical COVID-19 patients revealed an overexpression of NEU1 in infiltrated neutrophils. Moreover, Oseltamivir or Zanamivir treatment of whole blood cells from severe COVID-19 patients reduces host NEU-mediated shedding of cell surface sialic acid and neutrophil overactivation. These findings suggest that neuraminidase inhibitors can serve as host-directed interventions to dampen neutrophil dysfunction in severe infections.

**At a Glance:** In a severe systemic inflammatory response, such as sepsis and COVID-19, neutrophils play a central role in organ damage. Thus, finding new ways to inhibit the exacerbated response of these cells is greatly needed. Here, we demonstrate that *in vitro* treatment of whole blood with the viral neuraminidase inhibitors Oseltamivir or Zanamivir, inhibits the activity of human neuraminidases as well as the exacerbated response of neutrophils. In experimental models of severe sepsis, oseltamivir decreased neutrophil activation and increased the survival rate of mice. Moreover, Oseltamivir or Zanamivir *ex vivo* treatment of whole blood cells from severe COVID-19 patients rewire neutrophil function.

## INTRODUCTION

Neutrophils are key components of the immune response against pathogens (1). However, during severe acute infections, such as sepsis and COVID-19, overactivated neutrophils infiltrate vital organs and release molecules including proteases, reactive oxygen species (ROS), and neutrophil extracellular traps (NETs) (2, 3). While such inflammatory mediators are essential to the control of infection, they can also damage healthy cells (4). Therefore, the function of neutrophils must be regulated to efficiently clear microorganisms with minimal detrimental effects to the host.

A number of mechanisms controlling neutrophil activation have been described (5). For instance, the contents of sialic acid (Sia) have been demonstrated to regulate leukocyte activation to microbial stimuli (6). The dense array of Sia present in the glycocalyx of all mammalian cells makes this monosaccharide a central molecule for many cellular processes including: cell-cell interactions, signal transduction, and transendothelial migration (7). Neuraminidases (NEUs) are enzymes found in both pathogens and mammalian hosts, which hydrolyze Sia residues linked to galactose, N-acetylgalactosamine or polySia residues on glycoconjugates, thereby regulating many physiological and pathological responses (8). In human neutrophils, shedding of surface Sia by microbial-derived NEUs leads to cellular activation, ROS production, and release of NETs (9–12). Additionally, it has been demonstrated that LPS induces membrane-associated NEU activation in murine or human macrophages and dendritic cells (13). Upon LPS binding to TLR4, NEU activity was shown to regulate NF-κB induction in macrophages, suggesting a role for this enzyme during cellular activation (13). Also, in Gram-negative experimental sepsis, leukocyte dysfunction is mediated by NEU activity and associated with exacerbated inflammatory response and high mortality rates (14, 15). As previous studies have demonstrated that pathogen-derived NEU stimulate neutrophils, (9, 10, 16), we investigated whether endogenous host NEUs can be targeted to regulate neutrophil dysfunction observed in severe infections.

Here, we have identified host NEU activation as a positive regulator of microbial-induced human neutrophil overactivation. Additionally, we have employed antiviral NEU inhibitors, Oseltamivir and Zanamivir, to explore this pathway and found that these drugs fine-tune the neutrophil dysfunction observed in sepsis and COVID-19. Together, our results show that NEU is a potential target for the control of neutrophil dysfunction and present Oseltamivir or Zanamivir as adjunctive therapy for severe infections.

## METHODS

### Human blood samples

Blood samples were collected from severe COVID-19 (n=6) or convalescent COVID-19 (n=8) patients (25 to 89 yr old) admitted in the ICU or Research Center on Asthma and Airway Inflammation at the UFSC University Hospital. Blood samples from sex-matched healthy donors were used as controls. The research protocol was approved by the Institutional Review Board of the UFSC (CAAE #82815718.2.0000.0121, #36944620.5.1001.0121).

### Evaluation of neutrophil response

Whole blood were incubated in the presence or absence of Oseltamivir (100 µM, Sigma-Aldrich), Zanamivir (30 µM, Sigma-Aldrich), LPS (1 µg/mL, *E. coli* 0127:b8, Sigma-Aldrich), LPS plus Oseltamivir or LPS plus Zanamivir. Total leukocytes were used to evaluate the effect of isolated neuraminidase from *Clostridium perfringens* (CpNEU) on neutrophils. Leukocytes were incubated in the presence or absence of CpNEU (10 mU, Sigma-Aldrich), CpNEU plus Oseltamivir or CpNEU plus Zanamivir. Next, neutrophil activation, phagocytosis, bacterial killing, ROS production and NETs release were evaluated.

### Mice

All animal procedures followed the ARRIVE guidelines and the international principles for laboratory animal studies (17). Protocols were approved by the Animal Use Ethics Committee of UFSC (CEUA #8278290818). C57BL/6 and Swiss mice were used for peritonitis- or pneumonia-induced severe sepsis. Mice were randomly pretreated and/or post treated by *per oral* (PO, 12/12 hr) with saline or Oseltamivir phosphate (10 mg/kg, Eurofarma, Brazil). Additional details and methods are found in **Supplementary Methods**.

## RESULTS

### LPS-induced surface Sia shedding in human neutrophils is mediated by NEU activity

As activated NEUs hydrolyze Sia residues linked to underlying galactose glycoconjugates (7), we estimate Sia levels on neutrophils after their activation. LPS treatment of whole blood from healthy donors significantly reduces the binding of *Maackia amurensis* II (MAL-II, a lectin that binds selectively to α2-3-over α2-6-linked Sia (18)) on neutrophils when compared to untreated cells (**Supplementary Fig. 2A**). Next, cells were stained with an Fc-chimera of Siglec-9, a sialic acid-binding protein that recognizes Sia in α2-3 and α2-6 linkages (19). Similarly, binding of Siglec-9-Fc (**Supplementary Fig. 2B**) is decreased on neutrophils treated with LPS, confirming a reduction of neutrophil Sia residues likely due to LPS-induced NEU activity in these cells. To test this hypothesis, we measured NEU activity in human leukocytes using the NEU substrate 4-MU-NANA (13) and validated the assay using CpNEU (**Fig. 1A-B**). Both clinically available NEU inhibitors Oseltamivir and Zanamivir reduce CpNEU activity (**Fig. 1A-B**). LPS-induced NEU activity on leukocytes was also significantly inhibited by Oseltamivir or Zanamivir (**Fig. 1C-D**). Moreover, these NEU inhibitors prevent LPS- or CpNEU-mediated reduction of MAL-II (**Fig. 1E-F; K-L**) or *Sambucus nigra* (SNA) binding (SNA, a lectin that binds selectively to α2-6-over α2-3-linked Sia) (**Fig. 1G-H; M-N**). Desialylation unmasks galactose residues that are recognized by the lectin peanut agglutinin (PNA) (20). NEU inhibitors prevent LPS- or CpNEU-mediated the increase of the PNA binding on neutrophils surface (**Fig. 1I-J; O-P**). Together, these results show that LPS-induced host NEU activity decreases Sia content on neutrophils, which can be inhibited by Oseltamivir and Zanamivir.

**Figure 1.**
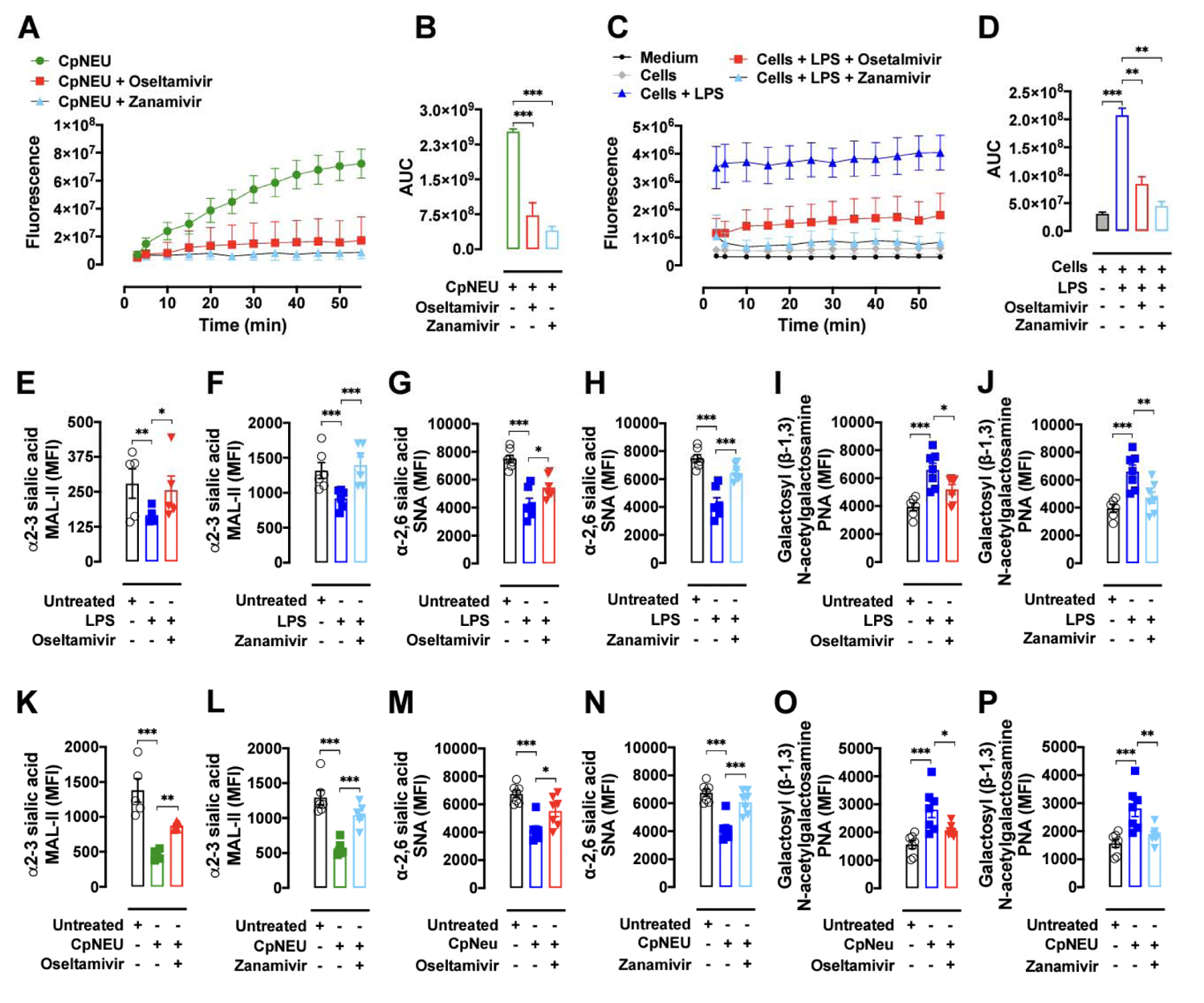
LPS stimulates NEU activity in human leukocytes. Neuraminidase isolated from *Clostridium perfringens* (CpNEU) was used to validate the NEU activity assay. CpNEU (0.012 UI) was added in a 96-well flat-bottom dark plate on ice in the presence or absence of Oseltamivir phosphate (100 µM) or Zanamivir (30 µM). Next, the substrate 4-MU-NANA (0.025 mM) was added and the fluorescent substrate was read 3 min after at 37 °C (**A**). The area under the curve (AUC) values are shown in **B**. Tota leukocytes resuspended in HBSS were added in a plate on ice and 4-MU-NANA substrate (0.025 mM) was added followed by the addition of medium, LPS (1 µg/mL), LPS plus Oseltamivir (100 µM) or LPS plus Zanamivir (30 µM). The fluorescent substrate was read 3 min after at 37 °C (**C**). Raw data were subtracted from the control group containing only HBSS (medium) and expressed as AUC values (**D**). Whole blood containing 1 x 10^6^ leukocytes from healthy donors were stimulated or not with LPS (1 µg/mL, 90 min, 37 °C, 5% CO_2_), LPS plus Oseltamivir (100 µM), or LPS plus Zanamivir (30 µM). Total leukocytes (1 x 10^6^) were incubated with CpNEU (10 mU, 60 min, 37 °C, 5% CO_2_), CpNEU plus Oseltamivir (100 µM), or CpNEU plus Zanamivir (30 µM). Leukocytes were stained with MAL-II to detect α2-3 sialic acids (**E-F; K-L**), with SNA to detect α2-6 sialic acids (**G-H; M-N**) or PNA to detect galactosyl (β-1,3) N-acetylgalactosamine (**I-J; O-P**). The MFI was analyzed on CD66b^+^/CD15^+^ cells using the gate strategies shown in Supplementary Fig. 1. \**P*< 0.05; \*\**P* < 0.01; \*\*\**P* < 0.001. This figure is representative of three independent experiments (n= 3-6) and data are shown as mean ± SEM. LPS = lipopolysaccharide; CpNEU = neuraminidase; MAL-II = *Maackia amurensis* lectin II; SNA = *Sambucus nigra* lectin; PNA = peanut agglutinin.

### LPS-induced phagocytosis and killing of *E. coli* is modulated by NEU activity

Bacteria uptake and killing are important functions of neutrophils (3). We next investigated whether host NEU regulates phagocytosis and killing of *E. coli.* Whole blood or total leukocytes from healthy donors were preincubated with LPS or CpNEU, respectively, and *E. coli* BioParticles® added to cells for 60 min. Ingested pHrodo *E. coli* by neutrophils were analyzed by flow cytometry. A significant increase in the MFI of unstimulated cells incubated with pHrodo *E. coli* was observed at 37 °C compared to cells at 4 °C (**Supplementary Fig. 3**). LPS (**Fig. 2A-C**) or CpNEU, used as a positive control of NEU effects (**Fig. 2D-F**), but not heat-inactivated CpNEU, significantly enhances phagocytosis of *E. coli*. Remarkably, these effects are inhibited by Zanamivir or Oseltamivir (**Fig. 2A-F**), suggesting that LPS-enhanced phagocytosis involves a host NEU-dependent pathway. Similarly, pretreatment of cells with LPS or CpNEU increases both the number of cells with bacteria as well as the number of bacteria per cell (**Fig. 2G-J**). These effects were also abolished when NEU inhibitors Oseltamivir and Zanamivir were added in the cell cultures (**Fig. 2G-J**). Furthermore, LPS or CpNEU treatment enhances intracellular and extracellular killing of *E. coli,* which are also inhibited by Oseltamivir or Zanamivir (**Fig. 2K-L**). These results suggest that NEU plays a critical role in LPS-upregulated phagocytosis and killing responses of neutrophils.

**Figure 2.**
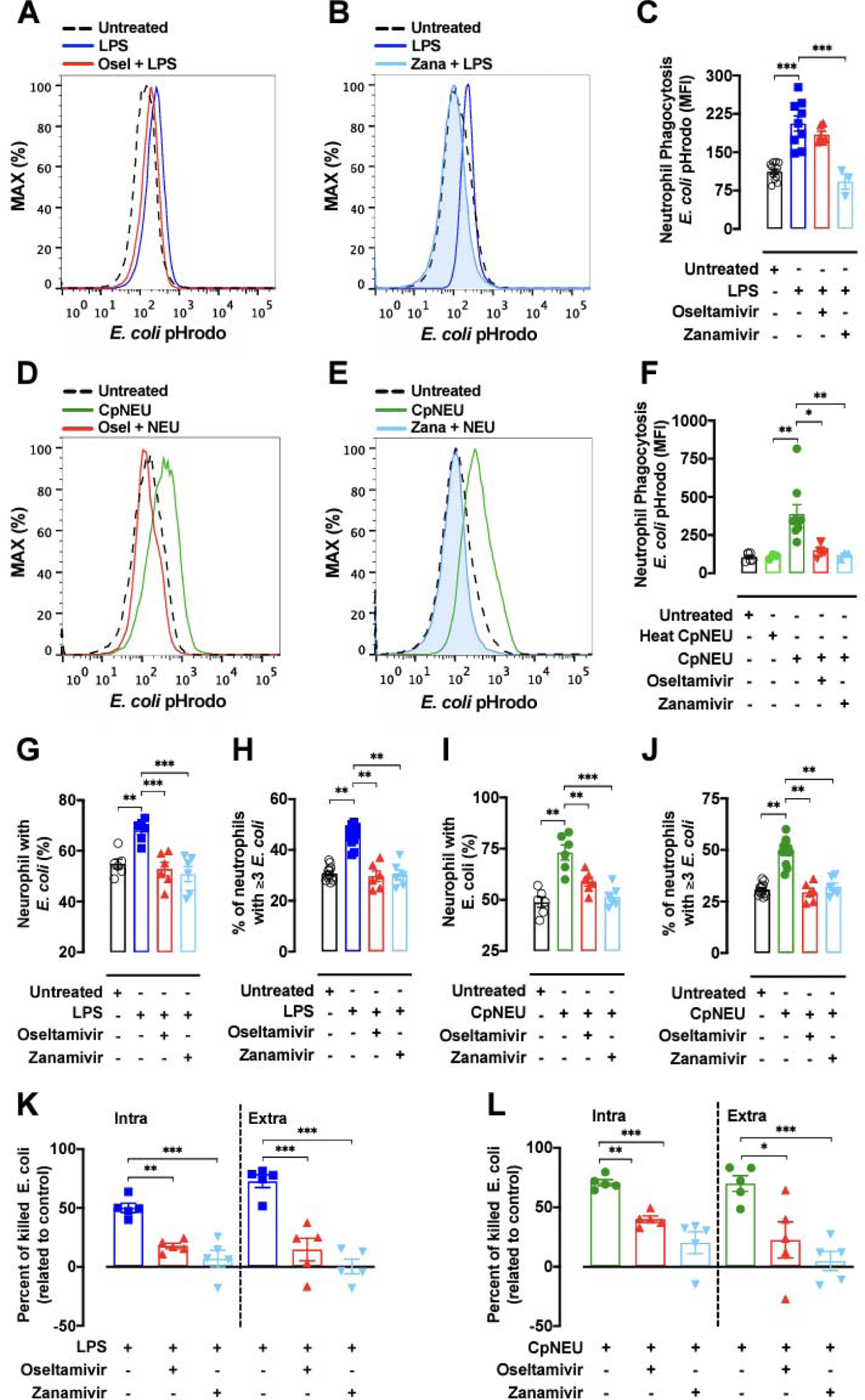
LPS increases phagocytosis and killing of *E. coli* in a NEU-dependent manner. Whole blood from healthy donors containing 1 x 10^6^ leukocytes were exposed (37 °C, 5% CO_2_) or not to LPS (1 µg/mL, 90 min), LPS plus Oseltamivir (100 µM), or LPS plus Zanamivir (30 µM) (**A-C; G-H; K**). Total leukocytes (1 x 10^6^) were exposed or not to CpNEU (10 mU, 60 min, 37 °C, 5% CO_2_), CpNEU plus Oseltamivir (100 µM), or CpNEU plus Zanamivir (30 µM) (**D-F; I-J; L**) and the phagocytosis and killing assays were performed. Leukocytes were incubated with *E. coli* pHrodo BioParticles® (100 µg/mL) for 60 min at 37 °C to assess phagocytosis in viable CD66b^+^/CD15^+^ cells (**A-F**) (as gated in Supplementary Fig. 1). Live *E. coli* was used to evaluate phagocytosis by light microscopy or to assess the killing by leukocytes. Cells were stimulated as described above and 1 x10^6^ leukocytes were incubated at 37 °C with *E. coli* (1 x10^6^ CFU) for 90 min for phagocytosis or for 180 min for killing assays. The percentage of cells with ingested bacteria (**G; I**) and the number of bacterial particles per cell (**H; J**, ≥3 particles per cell) were evaluated. The killing of *E. coli* was evaluated by spreading 10 µL of supernatant (extracellular killing) or 10 µL of the intracellular content in agar medium and the CFU were counted. Killing *E. coli* was expressed as the rate of fold change compared to the unprimed (untreated) cells (**L**). Symbols represent individual donors and data are shown as mean ± SEM from pooled data of two to three independent experiments (n = 3-12). \**P* < 0.05; \*\**P* < 0.01; \*\*\**P* < 0.001.

### NEU blockade prevents neutrophil activation

Shedding of cell surface Sia by mobilization of granule-associated NEU to the cell surface has been associated with neutrophil activation (21). Therefore, we analyzed surface expression of CD66b and CD62L, two markers of human neutrophil activation (22–24) and α2-3-Sia levels in LPS-exposed whole blood cultures. Both Oseltamivir and Zanamivir inhibit LPS-induced shedding of α2-3-Sia (**Fig. 3A,B**) and CD62L (**Fig. 3D,E**) or upregulation of CD66b (**Fig. 3G,H**) on neutrophils. Similarly, MAL-II preincubation, which prevents hydrolysis of α2-3-Sia by NEU due to steric hindrance at the enzyme cleavage site (25), blocks LPS-induced neutrophil activation (**Fig. 3C,F,I).** These data show that dampening NEU activity or blocking the hydrolysis of α2-3-Sia is sufficient to inhibit human neutrophil activation by LPS. Similar results were observed in soluble CpNEU-treated leukocytes (**Supplementary Fig. 4**). Next, we assessed whether NEU inhibitors influenced LPS-stimulated ROS production and release of NETs, key mediators of bacterial killing and tissue injury (26). Neutrophils primed with LPS and stimulated with PMA produce higher amounts of ROS when compared to unprimed cells (**Fig. 3J-L**). Both Oseltamivir and Zanamivir inhibit ROS release to levels similar to unprimed cells. These results were also reproduced by the treatment of cells with CpNEU (**Supplementary Fig. 5E-G**). Furthermore, Oseltamivir or Zanamivir significantly inhibit LPS-induced NETs released by isolated neutrophils (**Fig. 3M**). Together, these data indicate that microbial-induced host NEU activity regulates important neutrophil functions *in vitro*.

**Figure 3.**
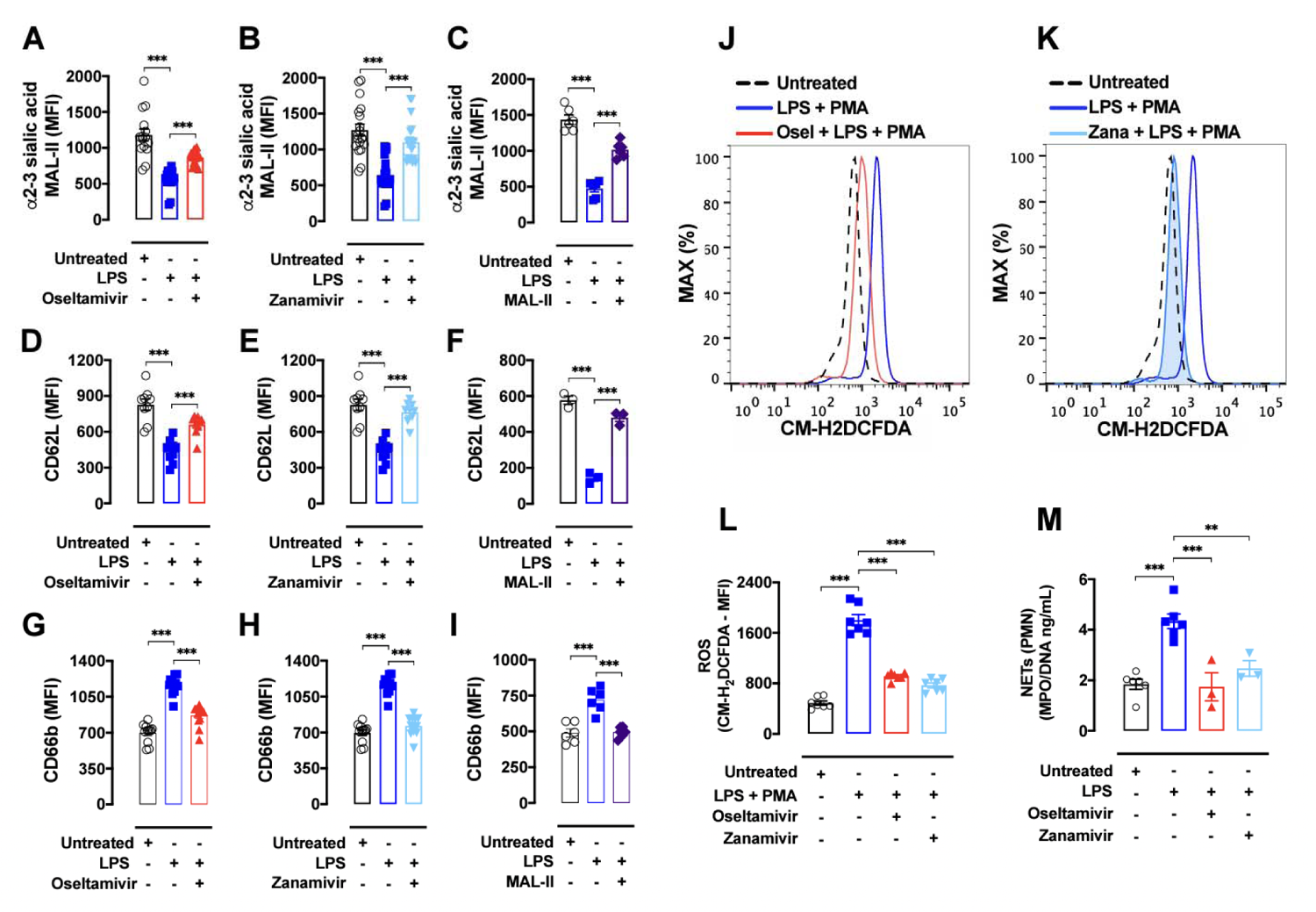
LPS-induced human neutrophil response involves NEU activity. Whole blood from healthy donors containing 1 x 10^6^ leukocytes were stimulated or not with LPS (1 µg/mL, 90 min, 37 °C, 5% CO_2_), LPS plus Oseltamivir (Osel, 100 µM), LPS plus Zanamivir (Zana, 30 µM), or LPS plus MAL-II (1 µg/mL, MAL-II promotes steric hindrance at the NEU cleavage site and prevent sialic acid cleavage). Leukocytes were marked with MAL-II (**A-C**) or with the cell activation markers CD62L (**D-F**) and CD66b (**G-I**). After red blood cells lysis leukocytes were incubated with 5 µM CM-H2DCFDA fluorescent probe for 15 min. PMA (10 µM) was used to stimulate ROS production for 10 min (**J-L**). Supplementary Fig. 5 showed ROS production in additional control groups. The MFI was analyzed on CD66b^+^ cells using the gate strategies shown in Supplementary Fig. 1. Isolated neutrophils were treated with Osetamivir (100 µM) or Zanamivir (30 µM) 1 h before the stimulus with LPS (10 µg/mL) for 4 h. The concentration of NETs was evaluated by MPO-DNA PicoGreen assay on supernatants of cells (**M**). Symbols represent individual donors and data are shown as mean ± SEM from pooled data of two to three independent experiments (n = 7) except for F and M that was made once with n=3. \*\*\**P* < 0.001; \*\**P* < 0.01. CM-H2DCFDA = 5-(and-6)-chloromethyl-2’,7’-dichlorodihydrofluorescein diacetate, acetyl ester; PMA = phorbol 12-myristate 13-acetate; ROS = reactive oxygen species; NETs = neutrophil extracellular traps; PMN = polymorphonuclear leukocytes.

### Oseltamivir enhances survival rate of mice in clinically relevant models of sepsis

Exacerbated neutrophil responses such as increased ROS production, release of NET and degranulation are associated with tissue injury and organ dysfunction (27). By using Oseltamivir as a therapeutic tool, we next explored the involvement of NEU activity *in vivo* during experimental sepsis, a model of neutrophil dysfunction (3, 28, 29). We first induced sepsis by intraperitoneal administration of 1 x 10^7^ CFU/mice of the Gram-negative *E. coli* (ATCC 25922), which lacks NEU in its genome (30). We used a dose of 10 mg/Kg of Oseltamivir by oral gavage, which is the equivalent dose used in humans (∼7.5 mg/Kg) (31). Oseltamivir pretreatment (2 hr before infection) plus post-treatment (6 hr after infection, 12/12 h, PO, for 4 days) markedly boost host survival (**Supplementary Fig. 6A**). Only a single dose of Oseltamivir 2 hr before bacterial administration significantly decreases the number of neutrophils in the BAL and lung tissue 4 or 6 hr after infection (**Supplementary Fig. 6B-C**). This pretreatment also augments neutrophil migration to the focus of infection, which is associated with an efficient control of infection (**Supplementary Fig. 6D-F**). Furthermore, pretreatment with Oseltamivir decreases BAL and plasma TNF and IL-17 levels (**Supplementary Fig. 6G-J**) and tissue injury markers (AST, ALT, ALP and total bilirubin) (**Supplementary Fig. 6K-N**), as well as prevents reduction of α2-3-Sia on peritoneal lavage SSC^high^/GR-1^high^ cells (**Supplementary Fig. 6O-P**). More importantly, the post-treatment efficacy of Oseltamivir was also evaluated in survival of septic mice. Mice were IP challenged with *E. coli* (1 x 10^7^ CFU/mice) and treated 6 hr after infection with Oseltamivir for 4 days (10 mg/Kg, PO, 12/12h). Strikingly, in the post-treatment protocol, Oseltamivir provides a significant improvement in the survival rate of septic mice (**Supplementary Fig. 6Q**).

Next, we employed the CLP model to evaluate the effect of Oseltamivir in septic mice, as it is considered the gold standard in preclinical sepsis (32). Six hours after CLP, Oseltamivir post-treatment leads to a small delay in the mortality rate of severe septic mice (**Supplementary Fig. 7A**). Next, CLP septic mice were treated with antibiotics because it is one of the standard interventions used in clinical settings of sepsis (33). Importantly, compared to the control animals, therapeutic use of Oseltamivir plus antibiotics drastically improved survival rates of CLP mice (87.5% experimental group vs 25% control group) (**Supplementary Fig. 7B**). Forty-eight hr after surgery, post-treated septic mice have a significant reduction of neutrophils in BAL and lungs, improvement of neutrophil migration at the focus of infection, and reduced bacterial load in PL and blood (**Supplementary Fig. 7C-G**). Levels of TNF and IL-17 in PL and plasma and tissue injury markers were also reduced in Oseltamivir treated mice (**Supplementary Fig. 7H-O**). Additionally, Oseltamivir also leads to a higher expression of α2-3-Sia on SSC^high^/GR-1^high^ cells in PL (**Supplementary Fig. 7P-Q**) confirming blockade of NEU activity *in vivo*.

As respiratory tract infections, particularly pneumonia, are among the most common sites of infection in sepsis (34), we intratracheally administered *K. pneumoniae* (ATCC 700603) into mice to address the effect of Oseltamivir. Post-treatment of mice with Oseltamivir significantly improves survival of septic mice challenged with *K. pneumoniae* (**FIG. 4A**). The increased host survival was accompanied by a decrease of neutrophil migration in BAL, reduced levels of TNF and IL-17 and reduced levels of tissue injury markers (**Fig. 4B-K**). Oseltamivir also prevents reduction of α2-3-Sia on BAL SSC^high^/GR-1^high^ cells (**Fig. 4L-M**). Together, these results show that host NEU activation exacerbates inflammatory responses during sepsis and the use of Oseltamivir improves disease outcome.

**Figure 4.**
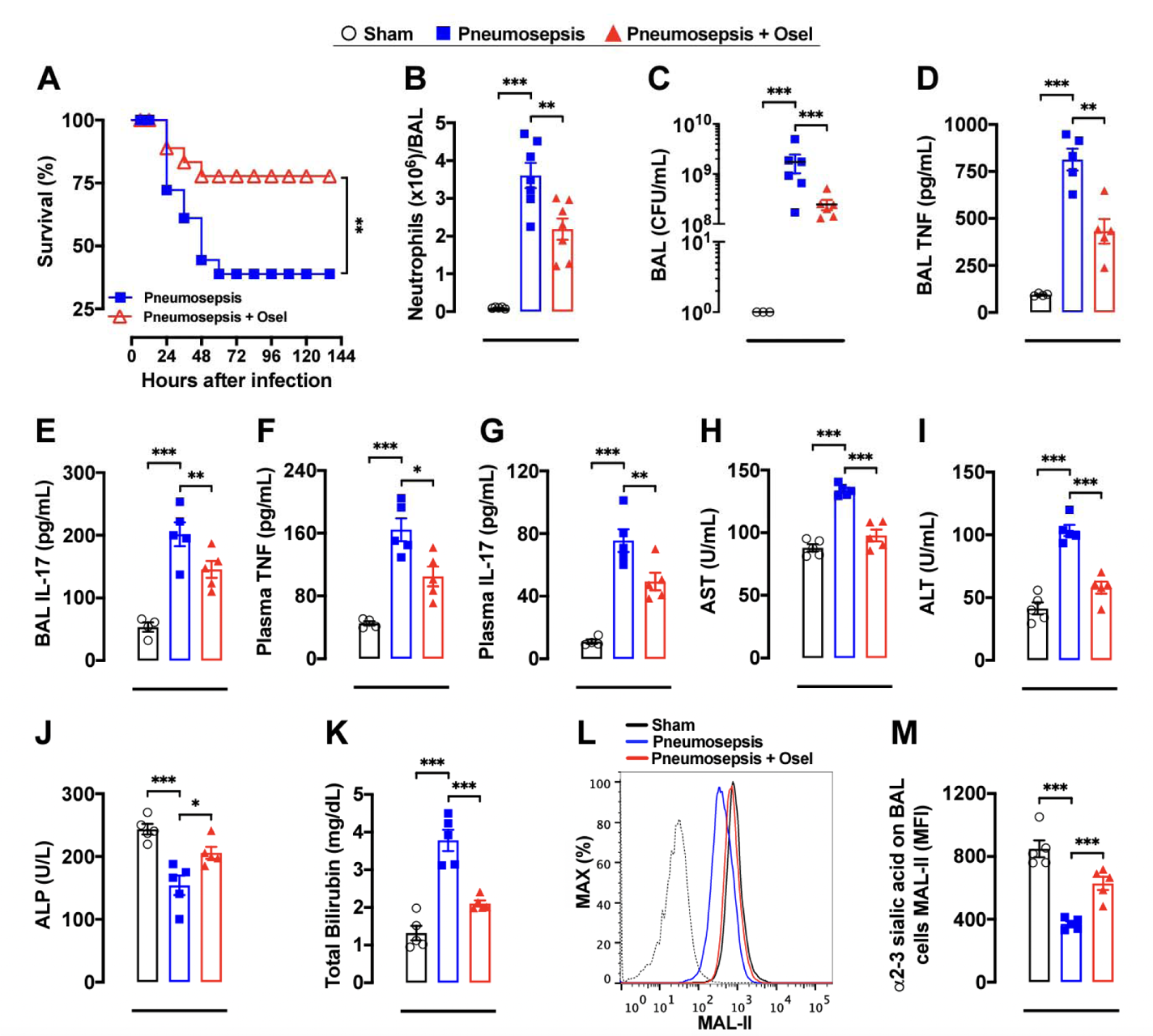
Oseltamivir enhanced mice survival in *K. pneumoniae*-induced sepsis. Sepsis was induced by intratracheal administration of *K. pneumoniae* and mice were randomly treated (starting 6 hr after infection, 12/12 hr, PO, n=20) with saline or Oseltamivir phosphate (10 mg/kg) and survival rates were monitored for 144 hr (**A**). In similar set of experiments, septic mice (n=6-7) were treated 6 hr after infection with a single dose of Oseltamivir phosphate (10 mg/kg, PO) and mice were euthanized 24 hr after infection to determine the number of neutrophils (**B**) and CFUs (**C**), and levels of TNF (**D**) and IL-17 (**E**) in BAL. Plasma levels of TNF (**F**), IL-17 (**G**), AST (**H**), ALT (**I**), ALP (**J**) and total bilirubin (**K**) were also evaluated 24 hr after infection. The amount of surface α2-3 sialic acids were assessed by MAL-II staining in SSC^high^/Gr-1^high^ cells in BAL and analyzed by FACS, as shown by the representative histograms (**L**) and MFI (**M**); dotted line = unstained cells. The results are expressed as percent of survival, mean or median (only for FACS data) ± SEM. \**P* < 0.05; \**P* < 0.01; \*\*\**P* < 0.001. Sham = sham-operated mice; Osel = Oseltamivir; AST = alanine aminotransferase; ALT = aspartate aminotransferase; ALP = alkaline phosphatase.

### Oseltamivir and Zanamivir rescue overactivated neutrophils from COVID-19 patients

Similar to bacterial sepsis, recent evidence suggests that neutrophils fuel hyper-inflammatory response during severe SARS-CoV-2 infection. Larger numbers of circulating neutrophils have been associated with poor prognosis of COVID-19 patients and analysis of lung biopsies and autopsy specimens showed extensive neutrophil infiltration (2, 35–41). Further studies by Chua *et al.* (2020) employing single-cell RNA sequencing (scRNA-seq) from upper and lower respiratory tract samples from COVID-19 patients indicated a significant augmentation of neutrophils linked to transcriptional program associated with tissue damage in epithelial and immune cells (42). Re-analysis of this scRNA-seq data (42) shows that expression of NEU1, but not NEU3 or NEU4, is upregulated in resident cells and highly expressed in neutrophils found in nasopharyngeal/pharyngeal swabs from COVID-19 patients (**Fig. 5**). A similar profile of NEUs expression was also observed in lower respiratory tract samples from these patients (**Supplementary Fig. 8**). These results suggest NEU enzymes are highly (41) expressed by lung infiltrating neutrophils during severe COVID-19. As demonstrated by Schulte-Schrepping *et al*. (2020) (43), circulating neutrophils are highly activated on active, but not convalescent, COVID-19 patients as observed by CD62L shedding (**Fig. 6A**) and upregulation of CD66b (**Fig. 6B**). Moreover, neutrophils from severe COVID-19 patients were found to present a significant reduction of surface α2-3-Sia (**Fig. 6C,D**), suggesting that NEU activity is also increased in blood neutrophils during severe COVID-19. Therefore, we asked whether neuraminidase inhibitors can rescue neutrophil activation from COVID-19 patients. *Ex vivo* treatment of whole blood with Oseltamivir or Zanamivir decreased neutrophil activation and restored the levels of cell surface sialic acid (**Fig. 6E-I**). As soluble NEU enzymes are also present in plasma (14), we next asked if plasma from COVID-19 patients can induce neutrophil response from healthy donors. Indeed, stimulation of whole blood from healthy donors with fresh plasma from severe, but not convalescent, COVID-19 patients leads to neutrophil activation (**Fig. 6J**), reduction of α2-3-Sia (**Fig. 6K**) as well as ROS production (**Fig. 6L,M**), which were significantly reduced by Oseltamivir or Zanamivir (**Fig. 6J-M**). Additionally, activity of NEU is increased in plasma from severe COVID-19 patients (**Supplementary Fig. 9A)**. Plasma samples from severe COVID-19 patients that were heat-inactivated to inhibit soluble NEU activity **(Supplementary Fig. 9B**) still induces neutrophil activation (**Fig. 6J-M**), suggesting that cellular NEU in conjunction with circulating factors mediate NEU-dependent neutrophil activation in severe COVID-19. These results highlight host NEU as a regulator of neutrophil activation in severe COVID-19 and suggest this pathway as a potential host-directed intervention target to rewire neutrophil responses during severe diseased.

**Figure 5.**
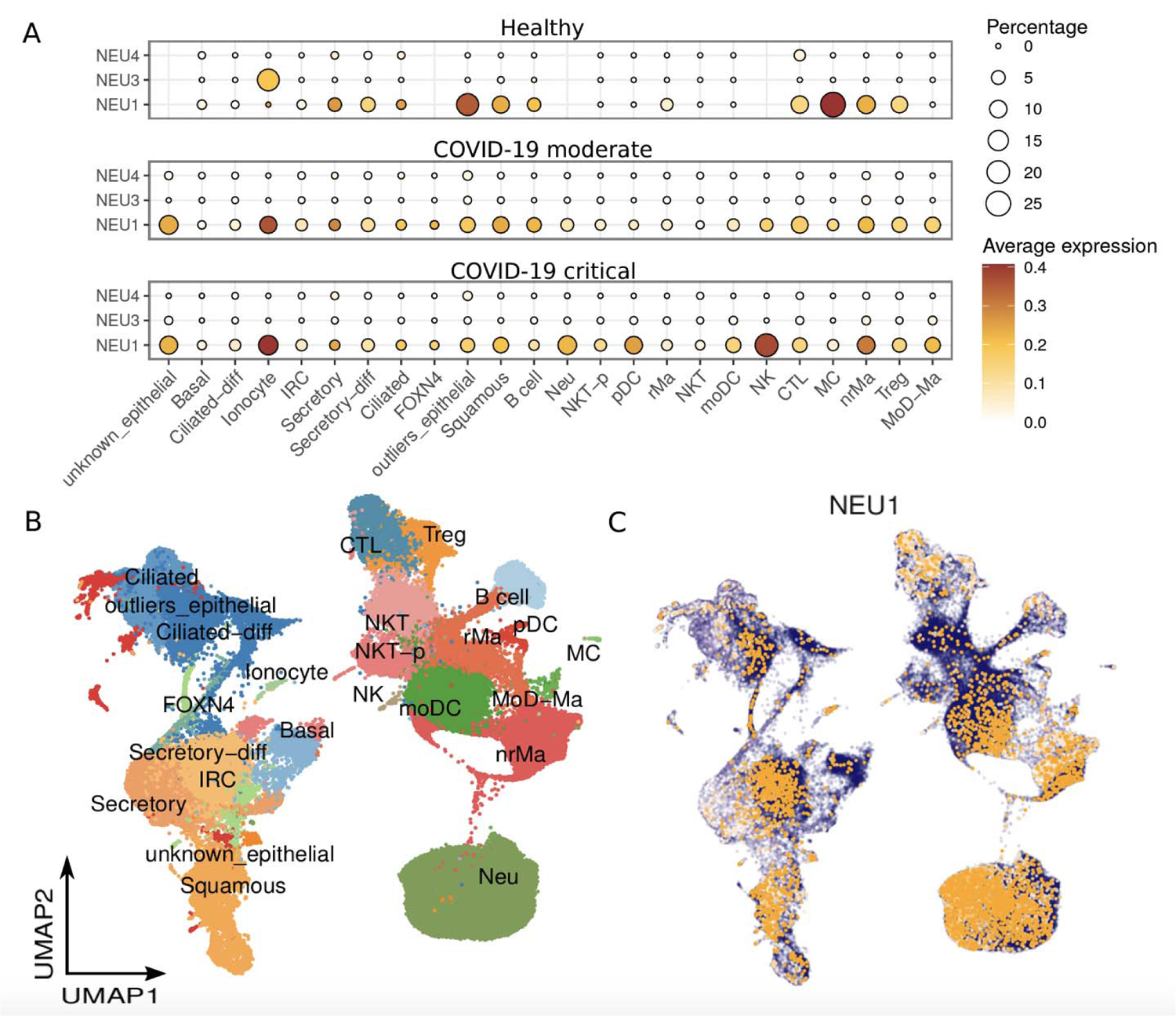
High expression of NEU1 in cell types from COVID-19 patients. **(A)** Gene expression of NEU1, NEU3 and NEU4 across cell types in healthy donors and moderate or critical COVID-19 patients. Size of the circle is proportional to the percentage of cells expressing the reported genes at a normalized expression leve higher than one. **(B)** UMAP analysis colored-coded by cell types in nasopharyngeal/pharyngeal swabs samples from healthy donors and COVID-19 patients. **(C)** Normalized expression of NEU1 overlaid on the UMAP spaces.

**Figure 6.**
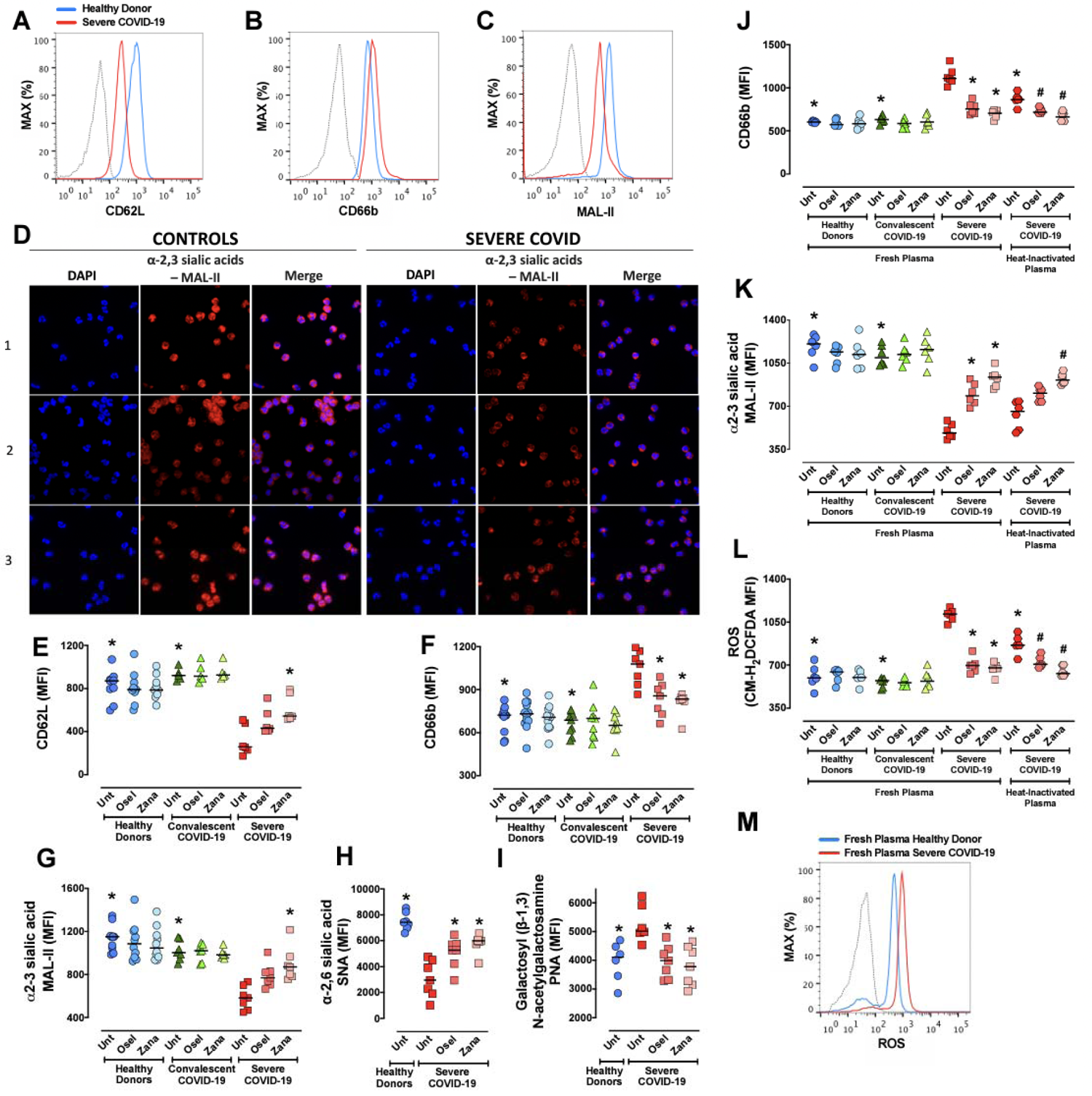
Oseltamivir and Zanamivir decrease neutrophil activation and increase sialic acid levels in active, but not convalescent neutrophils from COVID-19 patients. Whole blood from healthy donors (n= 10), severe COVID-19 patients (n= 6) or convalescent COVID-19 patients (n= 8) were treated or not with Oseltamivir (100 µM) or Zanamivir (30 µM) and total leukocytes were stained with the cell activation markers CD62L (**A and E**), CD66b (**B and F**) or the lectins MAL-II (**C, D and G)**, SNA (**H**) and PNA (**I**). Immunofluorescence (**D)** was carried out using biotinylated MAL-II followed by streptavidin Alexa Fluor 555 conjugate. Three different healthy donors (controls) and severe Covid-19 patients were used (magnification 100×). Blood samples from healthy donors (n = 7) were incubated for 2 h (37 °C, 5% CO_2_) with 7% of fresh plasma from healthy donors, severe or convalescent COVID-19 patients or with 7% of heat-inactivated plasma from severe COVID-19 patients in the presence or absence of Oseltamivir (100 µM) or Zanamivir (30 µM). Levels of CD66b (**J**), surface α2-3-Sia (MAL-II) (**K**), and ROS production (**L, M**) were assessed by FACS. The MFI was analyzed on CD66b^+^/CD15^+^ cells using the gate strategies shown in Supplementary Fig. 1. Symbols represent individual donors and data are shown as scatter dot plot with line at median from pooled data of two to seven independent experiments. The statistical significance between the groups was assessed by ANOVA followed by a multiple comparisons test of Tukey. The accepted level of significance for the test was P<0.05. * was significantly different when compared with Untreated Severe COVID-19; # was significantly different when compared with Untreated Heat-Inactivated Plasma from Severe COVID-19. Osel = Oseltamivir; Zana = Zanamivir.

## DISCUSSION

Systemic inflammatory responses may lead to unsuitable neutrophil stimulation, which is associated with higher mortality rates in sepsis and sepsis-like diseases (44). Therefore, finding new therapeutic options to prevent neutrophil overstimulation while maintaining their microbicidal abilities is greatly needed. Based on the findings presented here, NEU inhibitors are promising drugs to fill this gap. We demonstrated that endogenous host NEUs mediate exacerbated inflammatory responses by primary neutrophils. Clinically used viral NEU inhibitors, Oseltamivir and Zanamivir, decrease human NEU activity and are effective in preventing LPS-induced neutrophil responses or to rescue overactivation of neutrophils from COVID-19 patients. In severe murine sepsis, therapeutic use of Oseltamivir fine-tunes neutrophil migration promoting bacterial clearance and high survival rates.

All of the four different isotypes of NEU described in mammals (NEU1, NEU2, NEU3 and NEU4) remove Sia from glycoproteins and glycolipids with specific substrate preferences (45). NEU1 cleaves preferentially α2-3-Sia and seems to be the most important isoenzyme in immune cells. NEU1 is a lysosomal enzyme but it is also present at the cell surface where it can regulate multiple receptors such as Fc gamma receptor (FcγR), insulin receptor, integrin β-4, and TLRs (46). While several stimuli were described to induce NEU activity including LPS (13), PMA, calcium ionophore A23187, fMLP (21), and IL-8 (47), how NEUs are activated is poorly understood. However, NEU1 activation involves formation of a multicomplex of enzymes that stabilizes NEU1 in its conformational active state (48). Interestingly, NEU1 was found to be associated with matrix metalloproteinase-9 (MMP9) at the surface of naive macrophages (49). LPS binding to TLR4 leads to activation of a G protein-coupled receptor (GPCR) via GLi subunit and MMP9 to induce NEU1 activity, which in turn removes α2-3-Sia from TLR4, allowing its dimerization and intracellular signaling (25, 49, 50). Although we have not formally addressed whether the LPS-TLR4 pathway directly activates NEU function in human neutrophils, our results employing MAL-II preincubation suggest desialylation is required for LPS-mediated neutrophil responses. Thus, it is possible that NEU controls Sia levels in TLR4 molecules in human neutrophils as observed in macrophages and dendritic cells (25, 50).

The upstream involvement of NEU regulating LPS responses by neutrophils is in agreement with the previous demonstration that TLRs stimulate these cells independent of gene transcription (51). Together, our data suggests that NEU activation provides a fast response to enhance microbial-induced neutrophil functions. Thus, we speculate that this could be an evolutionary mechanism by which neutrophils quickly mobilize their microbicidal mediators against pathogens.

Sialic acid removal from the surface of neutrophils markedly changes their adhesiveness, chemotaxis, and migration (21, 47, 56–58). In peritonitis- or pneumonia-induced sepsis in mice, we observed that Oseltamivir prevented the massive neutrophil infiltration into bronchoalveolar spaces or lung tissues, suggesting that regulation of neutrophil migration by dampening NEU activity contributes to survival of septic mice. Interestingly, we observed a divergent effect of Oseltamivir on neutrophil migration to the focus of infection between peritonitis- and pneumonia-induced sepsis. This could be explained by the different mechanisms involved in neutrophil migration to the peritoneal cavity and lungs. While expression of CD62L and rolling of neutrophils to endothelium is necessary for its migration to the peritoneal cavity, it seems to be not required for migration into the lungs (3, 59). Moreover, systemic neutrophil activation leads to cell stiffening, resulting in retention of neutrophils in the small capillaries of the lungs (60), which is frequently the first organ impaired in non-pneumonia- and pneumonia-induced sepsis (61).

The role of NEU-induced neutrophil activation suggested here agrees with previous demonstration that NEU1 deletion in hematopoietic cells confers resistance to endotoxemia (15). Also, the sialidase inhibitor Neu5Gc2en protects endotoxemic irradiated wild-type (WT) mice reconstituted with WT bone marrow but not WT mice reconstituted with NEU1^-/-^ bone marrow cells (15). Similar to our findings, the treatment of mice with NEU inhibitors increases host survival in *E. coli*-induced sepsis (14). This outcome was correlated with significant inhibition of blood NEU activity. Enhancement of soluble NEU activity in serum decreases the Sia residues from alkaline phosphatase (APL) enzymes, which are involved in the clearance of circulating LPS-phosphate during sepsis (14).

SARS-CoV-2 infection leads to mild illness in most of the patients, but ∼20% of them progress to severe disease with many characteristics resembling sepsis, including acute respiratory distress syndrome (ARDS), cytokine storm, and neutrophil dysregulation (38, 62–64). The transcriptional programs found in neutrophil subsets from blood and lungs of severe COVID-19 patients are related to cell dysfunction, coagulation, and NETs formation (39, 43). We observed that blood neutrophils from severe COVID-19 are highly activated as demonstrated by reduced CD62L expression and increase of CD66b expression, as previously reported (43). We now add new information by showing that NEU1 is highly expressed in the respiratory tract of moderate and critical COVID-19 patients and blood neutrophils from severe, but not convalescent, COVID-19 patients with reduced surface levels of α2-3 and α2-6-Sia, suggesting a relevant role of NEU for COVID-19 pathology. More importantly, both the NEU inhibitors Oseltamivir and Zanamivir, increased the sialic acid content and rewired the overactivation of neutrophils from severe COVID-19 patients. We speculate that the addition of NEUs competitive inhibitors allowed the endogenous sialyltransferases to restore sialyl residues on surface glycoconjugates. Fast changes of surface sialic acid levels by sialidases and sialyltransferases seems to be an important mechanism to control neutrophil response (56). In neutrophils from healthy donors or COVID-19 convalescent patients, Oseltamivir and Zanamivir did not interfere in resting state and had no effect on sialic acid content, suggesting that NEU has a low effect on surface Sia turnover on non-activated neutrophils. How neutrophils are activated and the role of NEU in this process remains to be defined in COVID-19, nevertheless, recent evidence showed that neutrophils could be directly activated by SARS-CoV-2 (2), cytokines (65) and alarmins (39, 43), such as calprotectin (39), a TLR4 ligand (66). Furthermore, we now suggest that soluble NEU with other circulating factors present in plasma from severe COVID-19 patients also accounts for neutrophil activation.

Collectively, this work suggests that host NEU activation leads to shedding of surface sialic acid with consequent neutrophil overstimulation, tissue damage, and high mortality rates. On the other hand, NEU inhibitors-prevented shedding of sialic acid and regulates neutrophil response, resulting in infection control and high survival rates (working model in **Supplementary Fig. 10**). Considering that both drugs have a safety profile with few well-known adverse effects, our data suggest Oseltamivir and Zanamivir could be repurposed for the treatment of sepsis or severe infections such as COVID-19. Interestingly, a retrospective single-center cohort study including 1190 patients with COVID-19 in Wuhan, China, showed that administration of Oseltamivir was associated with a decreased risk of death in severe patients (67). Nevertheless, randomized clinical trials with NEU inhibitors in sepsis and COVID-19 are required to directly explore this hypothesis.

## Supporting information

Supplem methods and refererences

## Acknowledgments

We thank Universidade Federal de Santa Catarina (UFSC), the Laboratório Multiusuário de Estudos em Biologia (LAMEB/UFSC), CAPES/PrInt, CNPq and Le programme de bourses d’excellence Eiffel (Campus France - Ministère de l’Europe et des Affaires étrangères) for the support.

## Competing Interests statement

The authors declare that no conflict of interest exists.

This article has an online data supplement, which is accessible from this issue’s table of content online at www.atsjournals.org

## Supplementary Figure Legends

**Supplementary Fig. 1.**
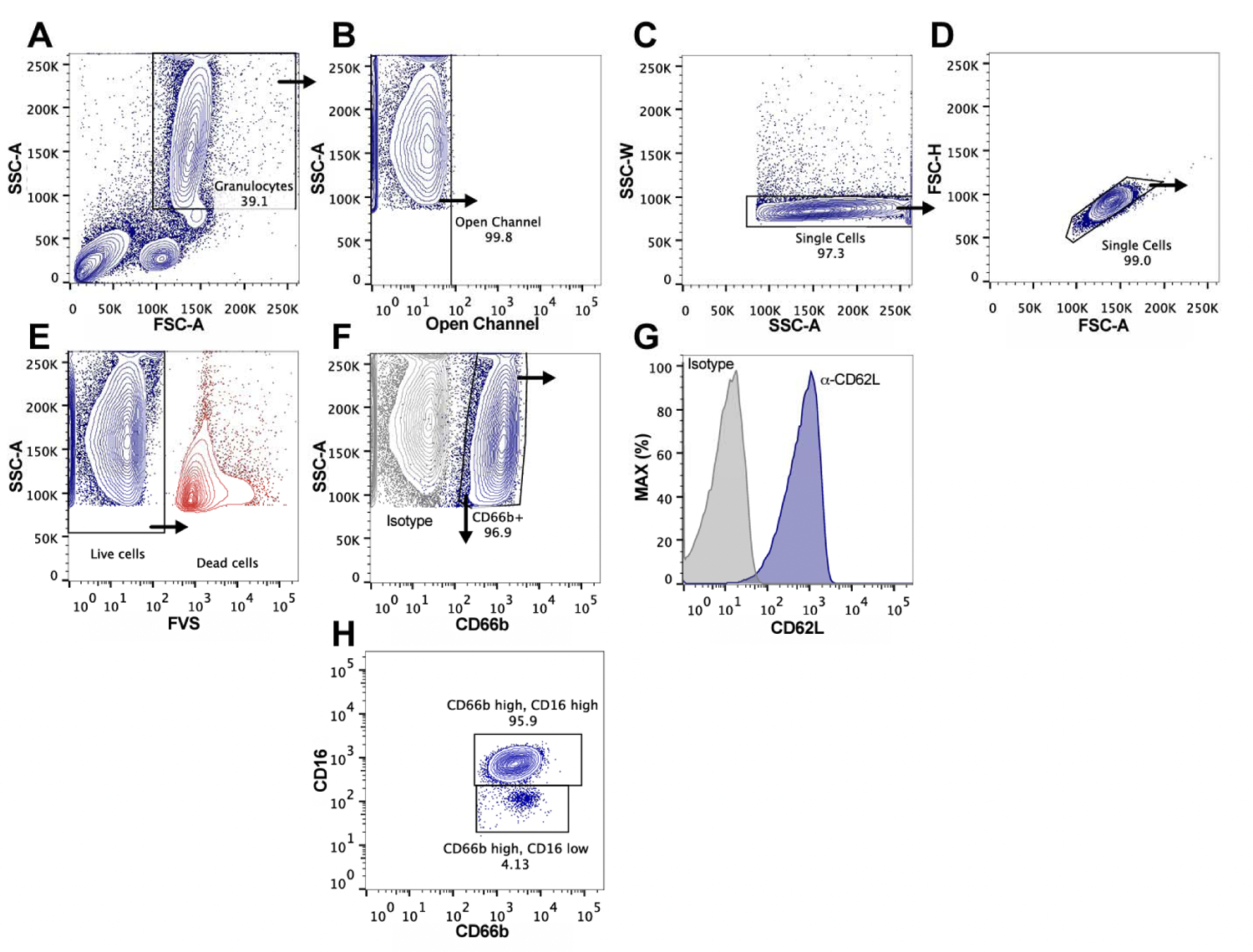
Representative gate strategy used for neutrophils analysis. A classic forward-scatter (FSC) vs side-scatter (SSC) characteristic dot plot was used to select neutrophils population (**A**) from peripheral blood collected from healthy donors and patients. Autofluorescent (**B**) and doublets (**C-D**) were excluded and live cells were selected (**E**). CD66b^+^ positive cells (**F**) were gated and the MFI of surface markers, such as CD62L (**G**) were assessed. Around 96% of CD66b^+^ cells are mature neutrophils (CD66b-high/CD16-high) and 4% of CD66b^+^ cells are CD66b-high/CD16-low, which is suggestive of immature neutrophils or eosinophils (**H**). Approximately 100.000 gated events were collected in each analysis. The analysis was performed in a FACSVerse using FACSuite software (BD Biosciences) and FlowJo software (FlowJo LLC).

**Supplementary Fig. 2.**
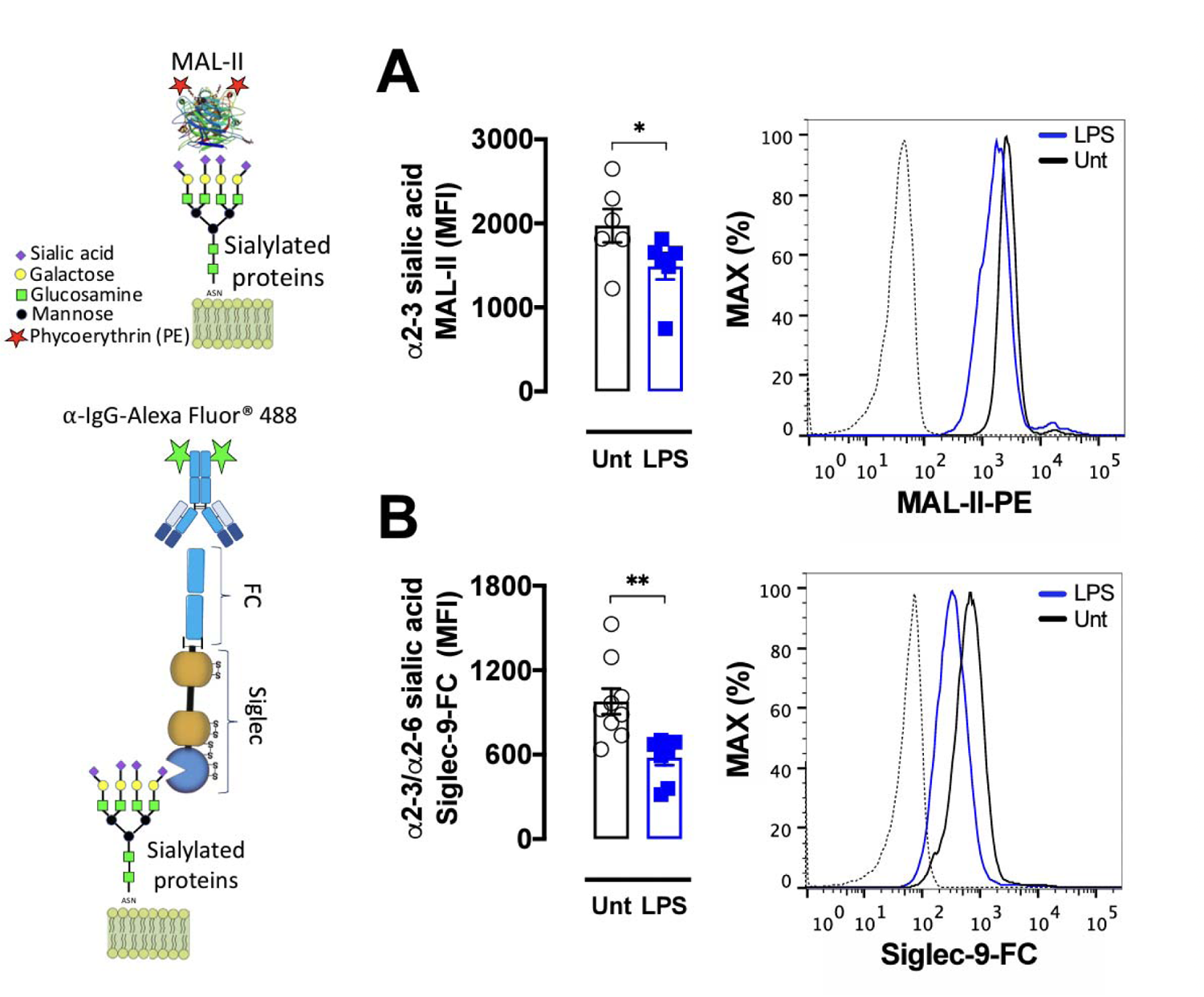
LPS reduces surface α2-3 sialic acids from human neutrophils. Whole blood containing 1 x 10^6^ leukocytes from healthy donors were stimulated with 1 µg/mL LPS for 90 min and α2-3 sialic acid contents were assessed by staining cells with biotinylated *Maackia Amurensis* Lectin II (MAL-II) (**A**) followed by streptavidin-phycoerythrin (PE) incubation. Siglec-9 ligands (**B**) were labeled by incubation of chimeric protein containing Siglec-9 sialic acid-Ig binding domain fused to a human IgG-Fc portion (Siglec-9-Fc). Siglec-Fc-9 were incubated with α-IgG1-Alexa Fluor 488 before adding the probe to cells. The MFI was analyzed on CD66b^+^ cells using the gate strategies shown in Supplementary Fig. 1. \**P* < 0.05. Symbols represent individual donors and data are shown as mean ± SEM from pooled data of two to three independent experiments (n=6-9). Unt = untreated cells; LPS = lipopolysaccharide; dotted line = unstained cells.

**Supplementary Fig. 3.**
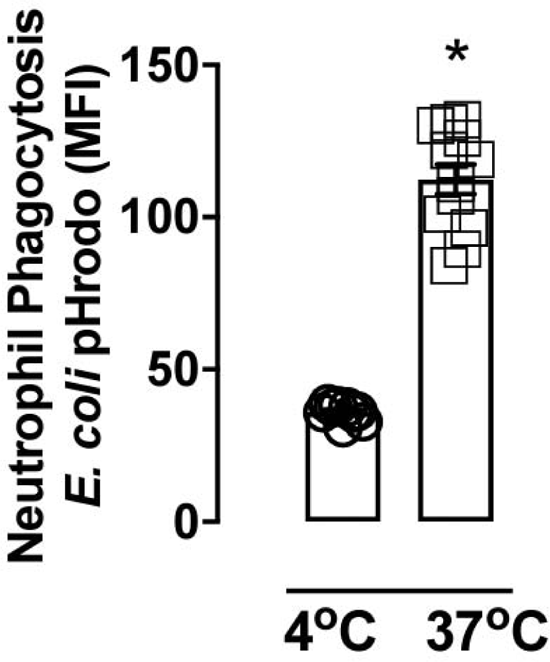
Phagocytosis of *E. coli* pHrodo bioparticles at 4 °C and 37 °C. Total leukocytes (1 x10^6^) were incubated with *E. coli* pHrodo bioparticles (100 µg/mL) for 60 min at 4°C or 37 °C and the phagocytosis in viable CD66b^+^ cells was assessed. Symbols represent individual donors and data are shown as mean ± SEM from pooled data of three to four independent experiments (n = 9-12). \**P* < 0.001.

**Supplementary Fig. 4.**
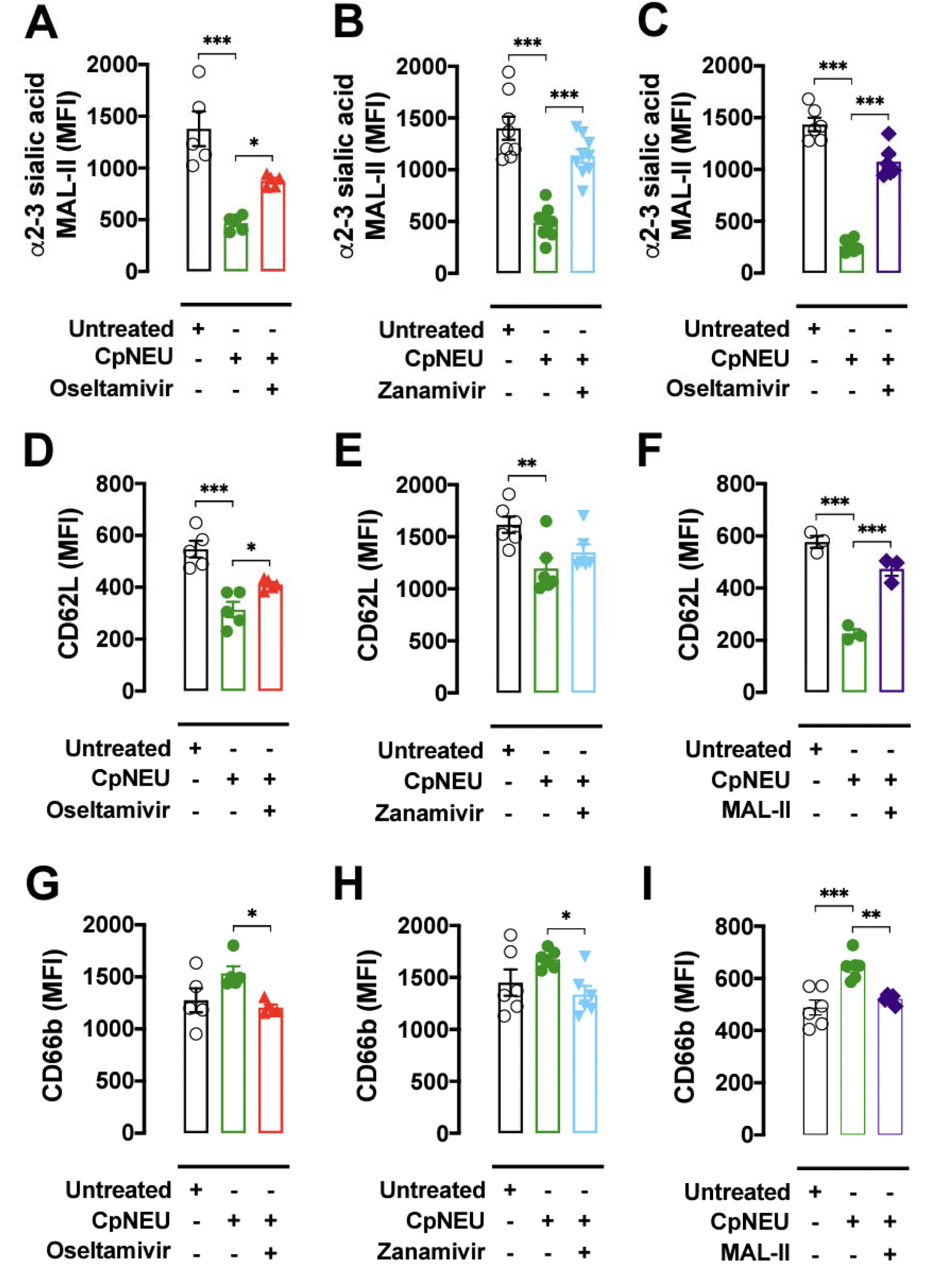
CpNeu-induced human neutrophil activation. Total leukocytes (1 x 10^6^) were incubated or not with CpNEU (10 mU, 60 min, 37 °C, 5% CO_2_) CpNEU plus Oseltamivir (100 µM), CpNEU plus Zanamivir (30 µM) or CpNEU plus MAL-II (1 µg/mL). Leukocytes were stained with MAL-II to detect α2-3 sialic acids (**A-C**) or with cell activation markers CD62L (**D-F**) and CD66b (**G-I**). The MFI was analyzed on CD66b^+^ cells. Symbols represent individual donors and data are shown as mean ± SEM from pooled data of two to three independent experiments (n = 5-9) except for F that was made once with n=3. \**P* < 0.05; \*\**P* < 0.01; \*\*\**P* < 0.001. MAL-II = *Maackia amurensis* lectin II; CpNEU = neuraminidase *Clostridium perfringens*.

**Supplementary Fig. 5.**
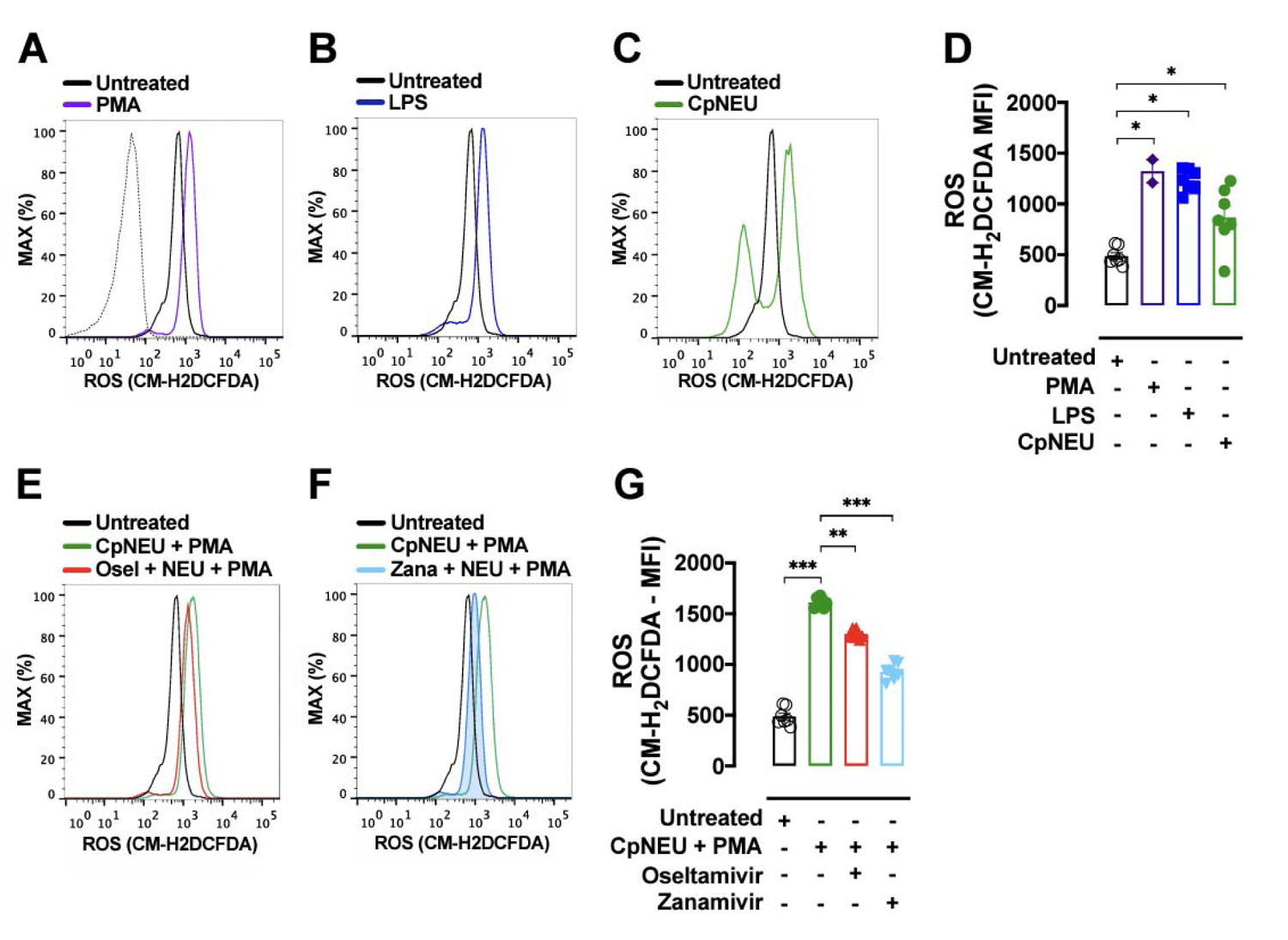
ROS production in neutrophils stimulated with LPS, CpNEU, or PMA. Whole blood from healthy donors containing 1 x 10^6^ leukocytes were exposed or not to LPS (1 µg/mL, 90 min) (**B and D**). Total leukocytes (1 x 10^6^) were incubated or not with CpNEU (10 mU, 60 min) (**C-D**), CpNEU plus Oseltamivir (100 µM) or CpNEU plus Zanamivir (30 µM) (**E-G**). Leukocytes were incubated with 5 µM CM-H2DCFDA fluorescent probe for 15 min and PMA (10 µM) was used to stimulate ROS production for 10 min (**A and E-G**). The MFI was analyzed on CD66b^+^ cells. Symbols represent individual donors and data are shown as mean ± SEM from pooled data of two independent experiments (n = 2-6). \**P* < 0.05; \*\**P* < 0.01; \*\*\**P* < 0.001. C = control; CM-H2DCFDA = 5-(and-6)-chloromethyl-2’,7’-dichlorodihydrofluorescein diacetate, acetyl ester; LPS = lipopolysaccharide; CpNEU = neuraminidase *Clostridium perfringens*; PMA = phorbol 12-myristate 13-acetate.

**Supplementary Fig. 6.**
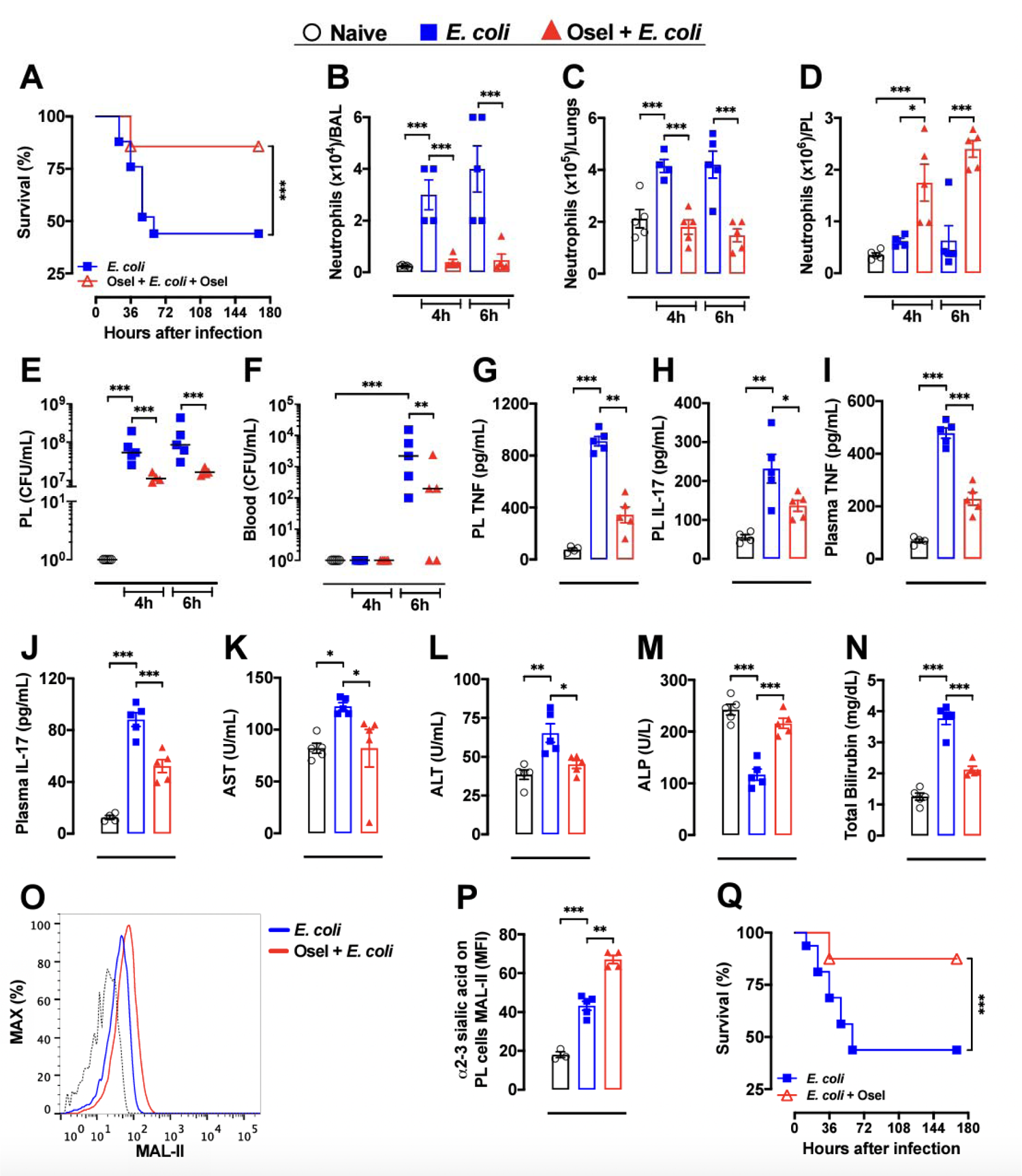
Oseltamivir improved the outcome of *E. coli*-induced sepsis. Sepsis was induced by intraperitoneal (IP) administration of 1 × 10^7^ CFU/mice *E. coli* (ATCC 25922). Mice were randomly pretreated *per oral* (PO) via (2 hr before infection) and posttreated (6 hr after infection, 12/12 hr, PO, for 4 days) with Oseltamivir phosphate (Osel, 10 mg/Kg) or saline and their survival rates were monitored over 168 hr (**A,** n=16). In another set of experiments (n=3-5) mice were randomly pretreated (2 hr before infection) with Oseltamivir phosphate (10 mg/Kg, PO) and the number of neutrophils in bronchoalveolar lavage (BAL, **B**) and in lung tissue (**C**) was counted. In peritoneal lavage (PL) infiltrating neutrophils counts (**D**), TNF (**G**), IL-17 (**H**) and the number of colony-forming units (CFU) in PL (**E**) or blood (**F**) were determined 4 or 6 hr after infection. Plasma levels of TNF (**I**), IL-17 (**J**), AST (**K**), ALT (**L**), ALP (**M**) and total bilirubin (**N**) were evaluated. The amount of surface α2-3 sialic acids were also assessed in PL SSC^high^/Gr-1^high^ cells as shown by the representative histograms (**O**) or MFI (**P**); dotted line = unstained cells. Mice were also randomly posttreated (starting 6 hr after infection, 12/12 hr, PO, for 4 days) with saline or Oseltamivir phosphate (10 mg/Kg) and their survival rates were monitored over 168 hr (**Q**). The results are expressed as percent of survival (n=16), mean or median (only for FACS data) ± SEM. \**P* < 0.05; \*\**P* < 0.01; \*\*\**P* < 0.001. These experiments were repeated 3 times for survival analysis and twice for other parameters. Osel = Oseltamivir; AST = alanine aminotransferase; ALT = aspartate aminotransferase; ALP = alkaline phosphatase; MAL-II = *Maackia amurensis* lectin II.

**Supplementary Fig. 7.**
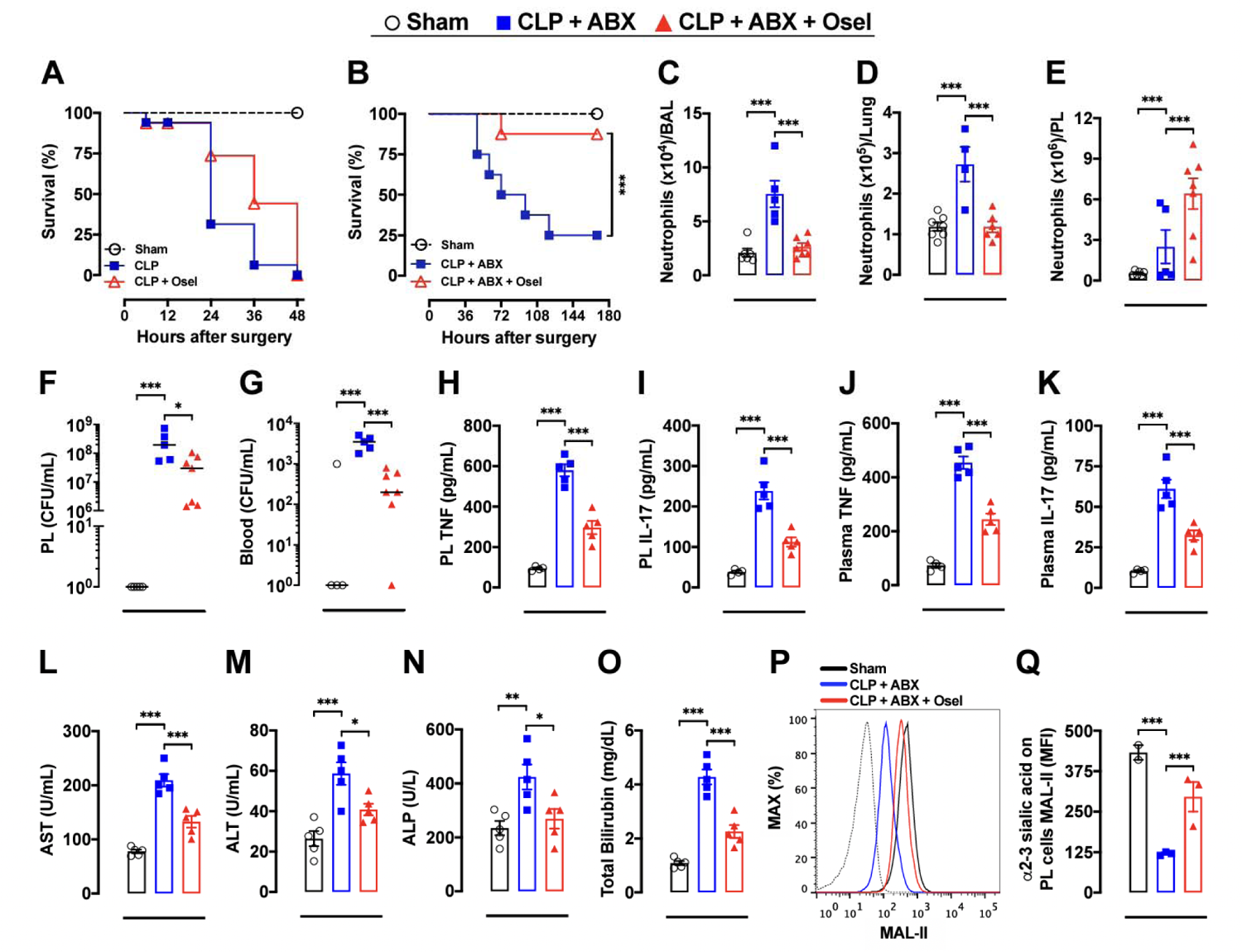
Oseltamivir enhanced host survival in CLP-induced sepsis. Severe sepsis was induced by the cecal ligation and puncture (CLP) model. Mice were randomly treated (starting 6 hr after infection, 12/12 h, PO, for 36 hr, n=16) with saline or Oseltamivir phosphate (10 mg/kg) and their survival rates were monitored over 48 hr (**A**). In another set of experiments, CLP mice were randomly IP treated (started 6 hr after infection, 12/12 hr) during 4 days with 100 µL metronidazole (15 mg/kg)/ceftriaxone (40 mg/kg) (ABX) plus saline or Oseltamivir phosphate (10 mg/kg) by PO and their survival rates (n=12) were monitored over 168 hr (**B**). Also, mice were subjected to CLP and treated with ABX + saline or ABX + Olsetamivir as described in B and euthanized 48 hr after surgery to evaluate the number of neutrophils in BAL (**C**), lung tissue (**D**), and peritoneal lavage (PL) (**E**); TNF (**H**), IL-17 (**I**), and CFU (**F**) were also determined in PL. Blood CFU (**G**) and plasmatic levels of TNF (**J**), IL-17 (**K**), AST (**L**), ALT (**M**), ALP (**N**) and total bilirubin (**O**) were also evaluated 48 hr after surgery. The amount of surface α2-3 sialic acids were assessed by MAL-II staining in SSC^high^/Gr-1^high^ cells in PL and analyzed by FACS, as shown by the representative histograms (**P**) and MFI (**Q**); dotted line = unstained cells. The results are expressed as percent of survival (n=16), mean or median (only for FACS data) ± SEM. \**P* < 0.05; \*\*\**P <* 0.001. These experiments were repeated 3 times for survival analysis and twice for other parameters (n=3-7). ABX = antibiotics (metronidazole/ceftriaxone); Sham = sham-operated. Osel = Oseltamivir; AST = alanine aminotransferase; ALT = aspartate aminotransferase; ALP = alkaline phosphatase; CFU = colony-forming units

**Supplementary Fig. 8.**
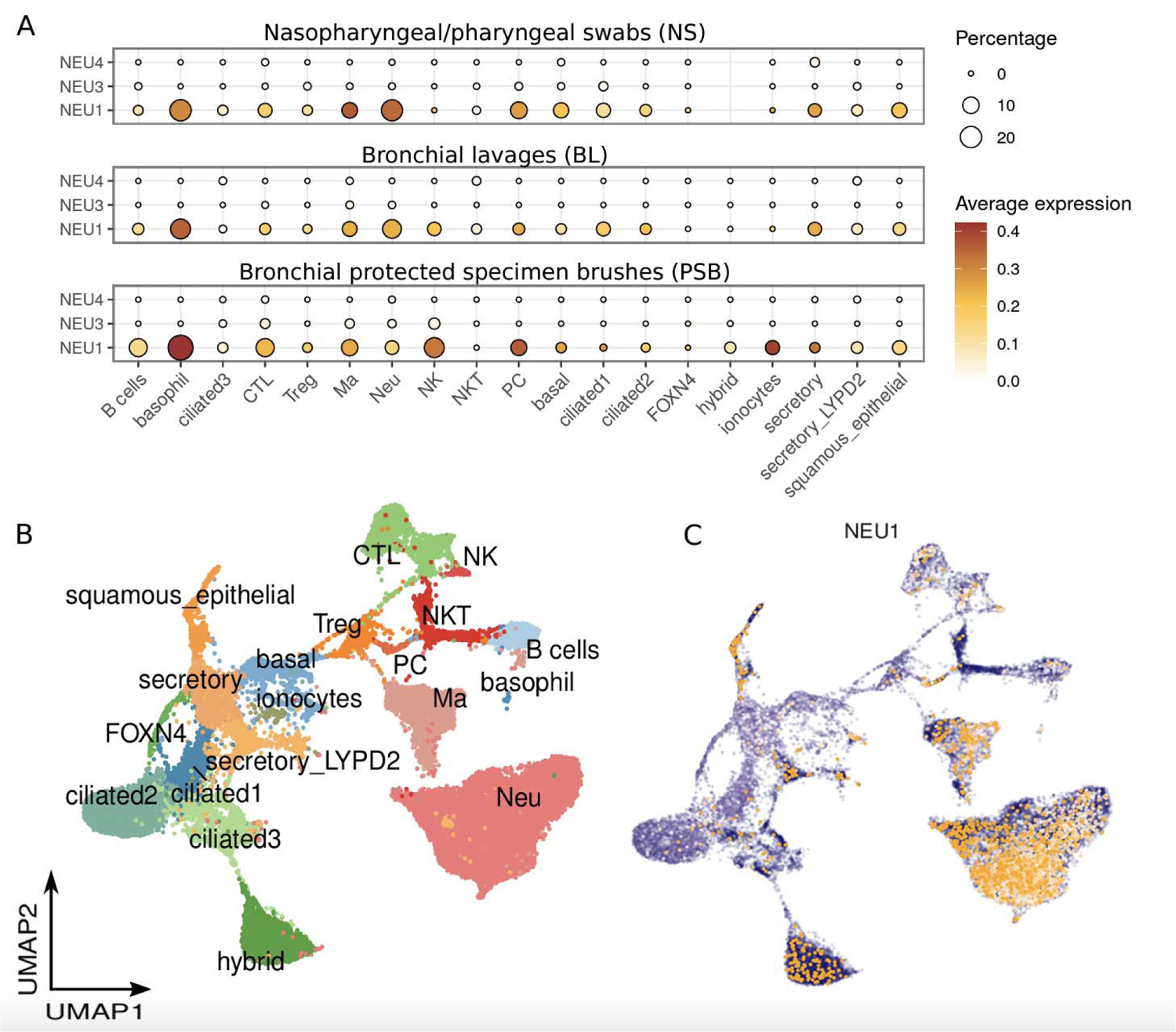
Expression of NEU1 in cell types from COVID-19 critical patients. **(A)** Gene expression of NEU1, NEU3 and NEU4 across cell types in two critical COVID-19 patients (BIH-CoV-01 and BIH-CoV-04). Size of the circle is proportional to the percentage of cells expressing the reported genes at a normalized expression level higher than one. **(B)** UMAP analysis colored-coded by cell types in NS, BL, and PSB samples from two critical COVID-19 patients. **(C)** Normalized expression of NEU1 overlaid on the UMAP space.

**Supplementary Fig. 9.**
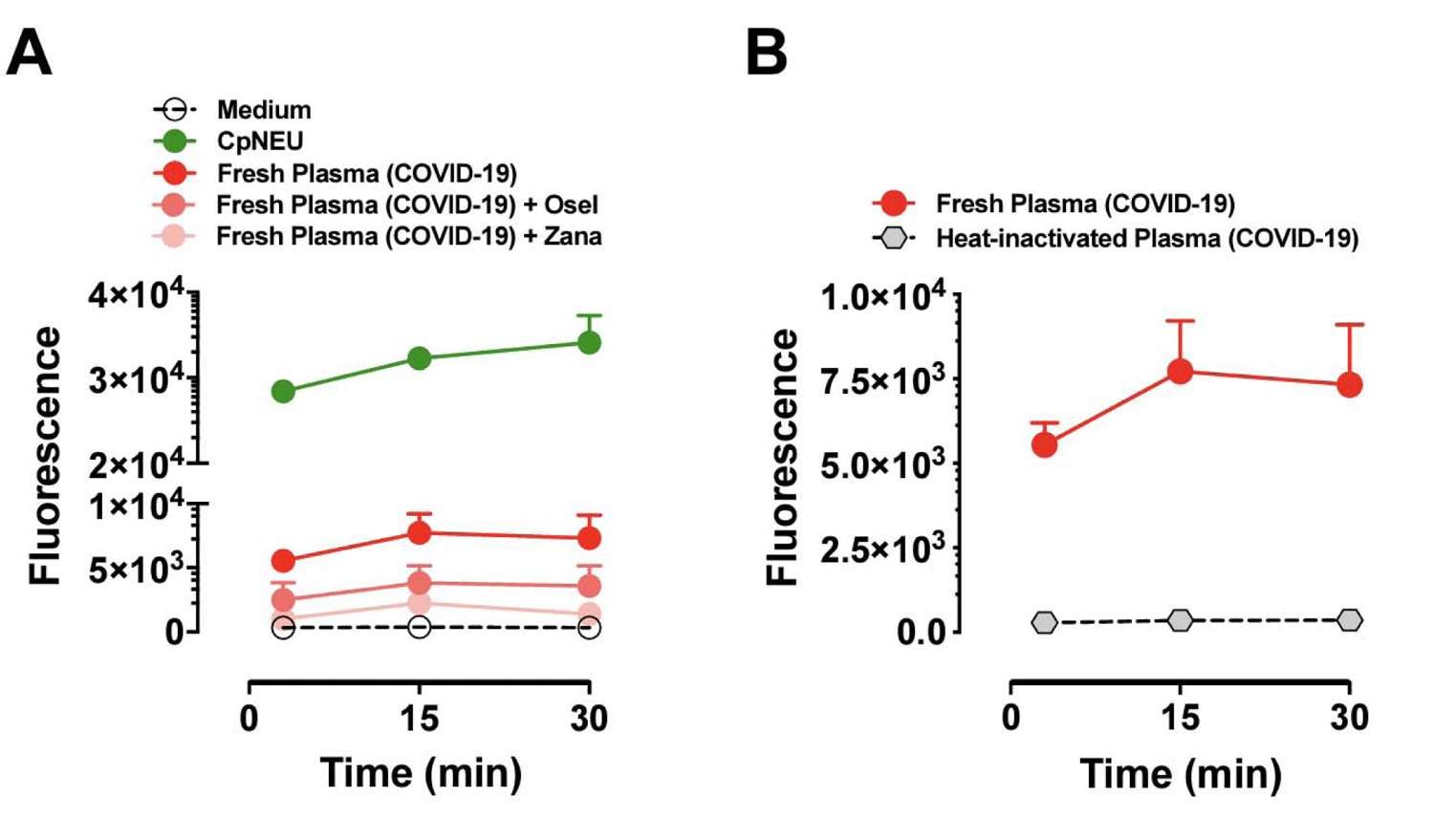
NEU activity is increased in plasma from severe COVID-19 patients. NEU activity was evaluated in fresh plasma from severe COVID-19 patients in the presence or absence of Oseltamivir (100 µM) or Zanamivir (30 µM) (**A**) and in heat-inactivated plasma from COVID-19 patients (**B**). Neuraminidase isolated from *Clostridium perfringens* (CpNEU) was used to validate the NEU activity assay. MAL-II = *Maackia amurensis* lectin II; CpNEU = neuraminidase *Clostridium perfringens*.

**Supplementary Fig. 10.**
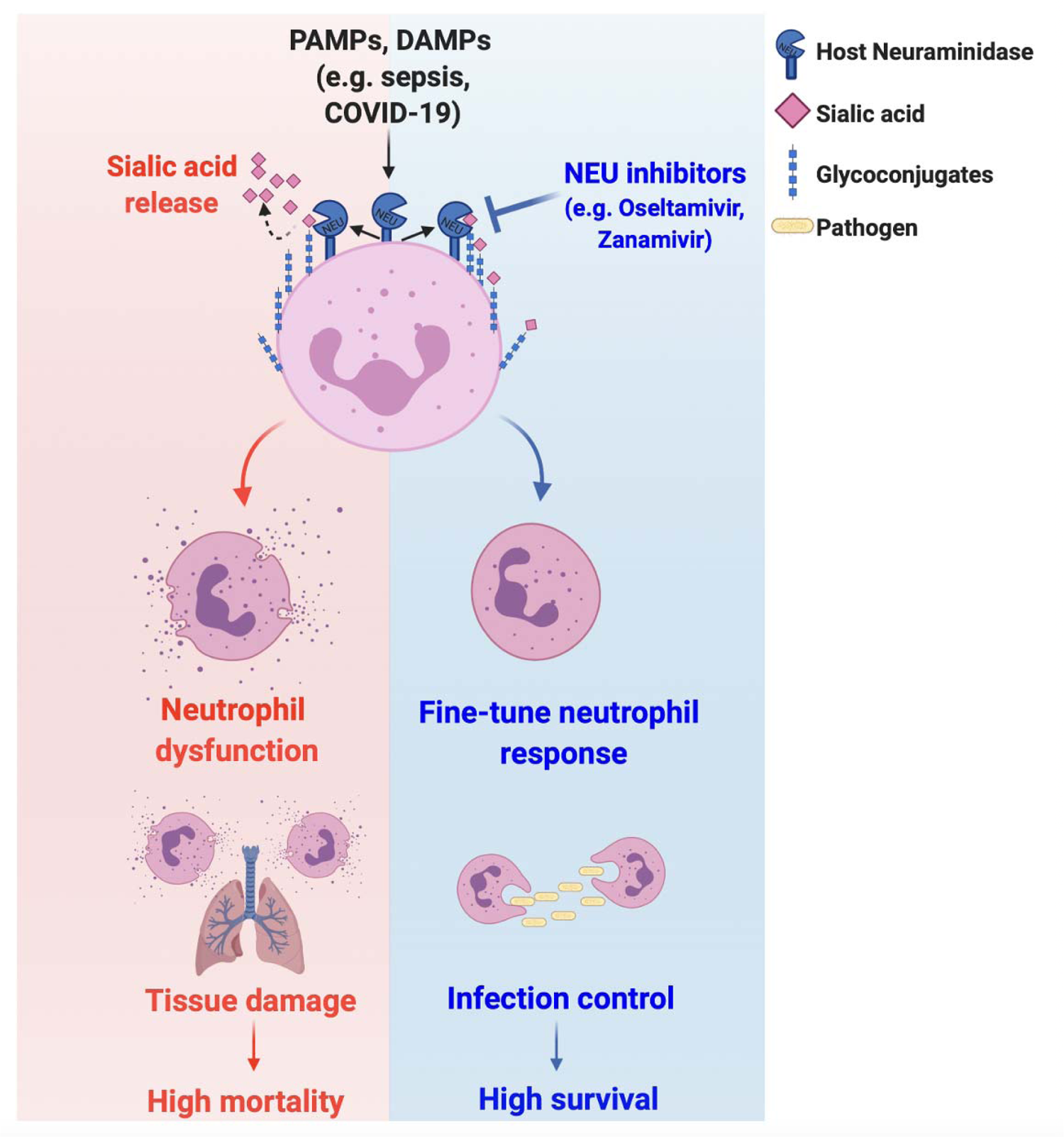
Working model. PAMPs and DAMPs in severe diseases such as sepsis and COVID-19 lead to neuraminidase activation with shedding of surface sialic acid and neutrophil overactivation, resulting in tissue damage and high mortality rates. On the other hand, neuraminidase inhibitors (e.g., Oseltamivir, Zanamivir) prevent the sialic acid release to regulate neutrophil response, resulting in infection control and high survival rates.

## REFERENCES

1. Mócsai A. Diverse novel functions of neutrophils in immunity, inflammation, and beyond. J Exp Med 2013;210:1283–1299.

2. Veras FP, Pontelli MC, Silva CM, Toller-Kawahisa JE, de Lima M, Nascimento DC, Schneider AH, Caetité D, Tavares LA, Paiva IM, Rosales R, Colón D, Martins R, Castro IA, Almeida GM, Lopes MIF, Benatti MN, Bonjorno LP, Giannini MC, Luppino-Assad R, Almeida SL, Vilar F, Santana R, Bollela VR, Auxiliadora-Martins M, Borges M, Miranda CH, Pazin-Filho A, da Silva LLP, et al. SARS-CoV-2-triggered neutrophil extracellular traps mediate COVID-19 pathology. J Exp Med 2020;217.:

3. Alves-Filho JC, Spiller F, Cunha FQ. Neutrophil paralysis in sepsis. Shock 2010;34 Suppl 1:15–21.

4. Segal AW. HOW NEUTROPHILS KILL MICROBES. Annual Review of Immunology 2005;

5. Steevels TAM, Meyaard L. Immune inhibitory receptors: essential regulators of phagocyte function. Eur J Immunol 2011;41:575–587.

6. Macauley MS, Crocker PR, Paulson JC. Siglec-mediated regulation of immune cell function in disease. Nat Rev Immunol 2014;14:653–666.

7. Varki A. Sialic acids in human health and disease. Trends in Molecular Medicine 2008;

8. Lipničanová S, Chmelová D, Ondrejovič M, Frecer V, Miertuš S. Diversity of sialidases found in the human body - A review. Int J Biol Macromol 2020;148:857–868.

9. Chang Y-C, Uchiyama S, Varki A, Nizet V. Leukocyte inflammatory responses provoked by pneumococcal sialidase. MBio 2012;3.:

10. Suzuki H, Kurita T, Kakinuma K. Effects of neuraminidase on O2 consumption and release of O2 and H2O2 from phagocytosing human polymorphonuclear leukocytes. Blood 1982;60:446–453.

11. Arora DJ, Henrichon M. Superoxide anion production in influenza protein-activated NADPH oxidase of human polymorphonuclear leukocytes. J Infect Dis 1994;169:1129–1133.

12. Henricks PA, van Erne-van der Tol ME, Verhoef J. Partial removal of sialic acid enhances phagocytosis and the generation of superoxide and chemiluminescence by polymorphonuclear leukocytes. J Immunol 1982;129:745–750.

13. Amith SR, Jayanth P, Franchuk S, Siddiqui S, Seyrantepe V, Gee K, Basta S, Beyaert R, Pshezhetsky AV, Szewczuk MR. Dependence of pathogen molecule-induced toll-like receptor activation and cell function on Neu1 sialidase. Glycoconj J 2009;26:1197–1212.

14. Yang WH, Heithoff DM, Aziz PV, Haslund-Gourley B, Westman JS, Narisawa S, Pinkerton AB, Millán JL, Nizet V, Mahan MJ, Marth JD. Accelerated Aging and Clearance of Host Anti-inflammatory Enzymes by Discrete Pathogens Fuels Sepsis. Cell Host Microbe 2018;24:500–513.e5.

15. Chen G-Y, Brown NK, Wu W, Khedri Z, Yu H, Chen X, van de Vlekkert D, D’Azzo A, Zheng P, Liu Y. Broad and direct interaction between TLR and Siglec families of pattern recognition receptors and its regulation by Neu1. Elife 2014;3:e04066.

16. Mills EL, Debets-Ossenkopp Y, Verbrugh HA, Verhoef J. Initiation of the respiratory burst of human neutrophils by influenza virus. Infection and Immunity 1981;

17. Kilkenny C, Browne WJ, Cuthill IC, Emerson M, Altman DG. Improving bioscience research reporting: the ARRIVE guidelines for reporting animal research. Osteoarthritis Cartilage 2012;20:256–260.

18. Knibbs RN, Goldstein IJ, Ratcliffe RM, Shibuya N. Characterization of the carbohydrate binding specificity of the leukoagglutinating lectin from Maackia amurensis. Comparison with other sialic acid-specific lectins. Journal of Biological Chemistry 1991;

19. Movsisyan LD, Macauley MS. Structural advances of Siglecs: insight into synthetic glycan ligands for immunomodulation. Org Biomol Chem 2020;18:5784–5797.

20. Varki A, Gagneux P. Multifarious roles of sialic acids in immunity. Ann N Y Acad Sci 2012;1253:16–36.

21. Cross AS, Wright DG. Mobilization of sialidase from intracellular stores to the surface of human neutrophils and its role in stimulated adhesion responses of these cells. J Clin Invest 1991;88:2067–2076.

22. Schmidt T, Zündorf J, Grüger T, Brandenburg K, Reiners A-L, Zinserling J, Schnitzler N. CD66b overexpression and homotypic aggregation of human peripheral blood neutrophils after activation by a gram-positive stimulus. J Leukoc Biol 2012;91:791–802.

23. Kishimoto TK, Jutila MA, Berg EL, Butcher EC. Neutrophil Mac-1 and MEL-14 adhesion proteins inversely regulated by chemotactic factors. Science 1989;245:1238–1241.

24. Jutila MA, Rott L, Berg EL, Butcher EC. Function and regulation of the neutrophil MEL-14 antigen in vivo: comparison with LFA-1 and MAC-1. J Immunol 1989;143:3318–3324.

25. Amith SR, Jayanth P, Franchuk S, Finlay T, Seyrantepe V, Beyaert R, Pshezhetsky AV, Szewczuk MR. Neu1 desialylation of sialyl alpha-2,3-linked beta-galactosyl residues of TOLL-like receptor 4 is essential for receptor activation and cellular signaling. Cell Signal 2010;22:314–324.

26. Mittal M, Siddiqui MR, Tran K, Reddy SP, Malik AB. Reactive oxygen species in inflammation and tissue injury. Antioxid Redox Signal 2014;20:1126–1167.

27. Sônego F, Castanheira FVES, Ferreira RG, Kanashiro A, Leite CAVG, Nascimento DC, Colón DF, Borges V de F, Alves-Filho JC, Cunha FQ. Paradoxical Roles of the Neutrophil in Sepsis: Protective and Deleterious. Front Immunol 2016;7:155.

28. Spiller F, Orrico MIL, Nascimento DC, Czaikoski PG, Souto FO, Alves-Filho JC, Freitas A, Carlos D, Montenegro MF, Neto AF, Ferreira SH, Rossi MA, Hothersall JS, Assreuy J, Cunha FQ. Hydrogen sulfide improves neutrophil migration and survival in sepsis via K+ATP channel activation. Am J Respir Crit Care Med 2010;182:360–368.

29. Spiller F, Carlos D, Souto FO, de Freitas A, Soares FS, Vieira SM, Paula FJA, Alves-Filho JC, Cunha FQ. α1-Acid glycoprotein decreases neutrophil migration and increases susceptibility to sepsis in diabetic mice. Diabetes 2012;61:1584–1591.

30. Vimr ER, Troy FA. Identification of an inducible catabolic system for sialic acids (nan) in Escherichia coli. J Bacteriol 1985;164:845–853.

31. Butler CC, van der Velden AW, Bongard E, Saville BR, Holmes J, Coenen S, Cook J, Francis NA, Lewis RJ, Godycki-Cwirko M, Llor C, Chlabicz S, Lionis C, Seifert B, Sundvall P-D, Colliers A, Aabenhus R, Bjerrum L, Jonassen Harbin N, Lindbæk M, Glinz D, Bucher HC, Kovács B, Radzeviciene Jurgute R, Touboul Lundgren P, Little P, Murphy AW, De Sutter A, Openshaw P, et al. Oseltamivir plus usual care versus usual care for influenza-like illness in primary care: an open-label, pragmatic, randomised controlled trial. Lancet 2020;395:42–52.

32. Rittirsch D, Huber-Lang MS, Flierl MA, Ward PA. Immunodesign of experimental sepsis by cecal ligation and puncture. Nat Protoc 2009;4:31–36.

33. Rhodes A, Evans LE, Alhazzani W, Levy MM, Antonelli M, Ferrer R, Kumar A, Sevransky JE, Sprung CL, Nunnally ME, Rochwerg B, Rubenfeld GD, Angus DC, Annane D, Beale RJ, Bellinghan GJ, Bernard GR, Chiche J-D, Coopersmith C, De Backer DP, French CJ, Fujishima S, Gerlach H, Hidalgo JL, Hollenberg SM, Jones AE, Karnad DR, Kleinpell RM, Koh Y, et al. Surviving Sepsis Campaign: International Guidelines for Management of Sepsis and Septic Shock: 2016. Intensive Care Med 2017;43:304–377.

34. Chou EH, Mann S, Hsu T-C, Hsu W-T, Liu CC-Y, Bhakta T, Hassani DM, Lee C-C. Incidence, trends, and outcomes of infection sites among hospitalizations of sepsis: A nationwide study. PLoS One 2020;15:e0227752.

35. Middleton EA, He X-Y, Denorme F, Campbell RA, Ng D, Salvatore SP, Mostyka M, Baxter-Stoltzfus A, Borczuk AC, Loda M, Cody MJ, Manne BK, Portier I, Harris ES, Petrey AC, Beswick EJ, Caulin AF, Iovino A, Abegglen LM, Weyrich AS, Rondina MT, Egeblad M, Schiffman JD, Yost CC. Neutrophil extracellular traps contribute to immunothrombosis in COVID-19 acute respiratory distress syndrome. Blood 2020;136:1169–1179.

36. Schurink B, Roos E, Radonic T, Barbe E, Bouman CSC, de Boer HH, de Bree GJ, Bulle EB, Aronica EM, Florquin S, Fronczek J, Heunks LMA, de Jong MD, Guo L, du Long R, Lutter R, Molenaar PCG, Neefjes-Borst EA, Niessen HWM, van Noesel CJM, Roelofs JJTH, Snijder EJ, Soer EC, Verheij J, Vlaar APJ, Vos W, van der Wel NN, van der Wal AC, van der Valk P, et al. Viral presence and immunopathology in patients with lethal COVID-19: a prospective autopsy cohort study. Lancet Microbe 2020;1:e290–e299.

37. Huang C, Wang Y, Li X, Ren L, Zhao J, Hu Y, Zhang L, Fan G, Xu J, Gu X, Cheng Z, Yu T, Xia J, Wei Y, Wu W, Xie X, Yin W, Li H, Liu M, Xiao Y, Gao H, Guo L, Xie J, Wang G, Jiang R, Gao Z, Jin Q, Wang J, Cao B. Clinical features of patients infected with 2019 novel coronavirus in Wuhan, China. Lancet 2020;395:497–506.

38. Guan W-J, Ni Z-Y, Hu Y, Liang W-H, Ou C-Q, He J-X, Liu L, Shan H, Lei C-L, Hui DSC, Du B, Li L-J, Zeng G, Yuen K-Y, Chen R-C, Tang C-L, Wang T, Chen P-Y, Xiang J, Li S-Y, Wang J-L, Liang Z-J, Peng Y-X, Wei L, Liu Y, Hu Y-H, Peng P, Wang J-M, Liu J-Y, et al. Clinical Characteristics of Coronavirus Disease 2019 in China. N Engl J Med 2020;382:1708–1720.

39. Silvin A, Chapuis N, Dunsmore G, Goubet A-G, Dubuisson A, Derosa L, Almire C, Hénon C, Kosmider O, Droin N, Rameau P, Catelain C, Alfaro A, Dussiau C, Friedrich C, Sourdeau E, Marin N, Szwebel T-A, Cantin D, Mouthon L, Borderie D, Deloger M, Bredel D, Mouraud S, Drubay D, Andrieu M, Lhonneur A-S, Saada V, Stoclin A, et al. Elevated Calprotectin and Abnormal Myeloid Cell Subsets Discriminate Severe from Mild COVID-19. Cell 2020;182:1401–1418.e18.

40. Wilk AJ, Rustagi A, Zhao NQ, Roque J, Martínez-Colón GJ, McKechnie JL, Ivison GT, Ranganath T, Vergara R, Hollis T, Simpson LJ, Grant P, Subramanian A, Rogers AJ, Blish CA. A single-cell atlas of the peripheral immune response in patients with severe COVID-19. Nat Med 2020;26:1070–1076.

41. Kuri-Cervantes L, Pampena MB, Meng W, Rosenfeld AM, Ittner CAG, Weisman AR, Agyekum RS, Mathew D, Baxter AE, Vella LA, Kuthuru O, Apostolidis SA, Bershaw L, Dougherty J, Greenplate AR, Pattekar A, Kim J, Han N, Gouma S, Weirick ME, Arevalo CP, Bolton MJ, Goodwin EC, Anderson EM, Hensley SE, Jones TK, Mangalmurti NS, Luning Prak ET, Wherry EJ, et al. Comprehensive mapping of immune perturbations associated with severe COVID-19. Sci Immunol 2020;5.:

42. Chua RL, Lukassen S, Trump S, Hennig BP, Wendisch D, Pott F, Debnath O, Thürmann L, Kurth F, Völker MT, Kazmierski J, Timmermann B, Twardziok S, Schneider S, Machleidt F, Müller-Redetzky H, Maier M, Krannich A, Schmidt S, Balzer F, Liebig J, Loske J, Suttorp N, Eils J, Ishaque N, Liebert UG, von Kalle C, Hocke A, Witzenrath M, et al. COVID-19 severity correlates with airway epithelium-immune cell interactions identified by single-cell analysis. Nat Biotechnol 2020;38:970–979.

43. Schulte-Schrepping J, Reusch N, Paclik D, Baßler K, Schlickeiser S, Zhang B, Krämer B, Krammer T, Brumhard S, Bonaguro L, De Domenico E, Wendisch D, Grasshoff M, Kapellos TS, Beckstette M, Pecht T, Saglam A, Dietrich O, Mei HE, Schulz AR, Conrad C, Kunkel D, Vafadarnejad E, Xu C-J, Horne A, Herbert M, Drews A, Thibeault C, Pfeiffer M, et al. Severe COVID-19 Is Marked by a Dysregulated Myeloid Cell Compartment. Cell 2020;182:1419– 1440.e23.

44. Leliefeld PHC, Wessels CM, Leenen LPH, Koenderman L, Pillay J. The role of neutrophils in immune dysfunction during severe inflammation. Crit Care 2016;20:73.

45. Smutova V, Albohy A, Pan X, Korchagina E, Miyagi T, Bovin N, Cairo CW, Pshezhetsky AV. Structural basis for substrate specificity of mammalian neuraminidases. PLoS One 2014;9:e106320.

46. Glanz VY, Myasoedova VA, Grechko AV, Orekhov AN. Sialidase activity in human pathologies. Eur J Pharmacol 2019;842:345–350.

47. Cross AS, Sakarya S, Rifat S, Held TK, Drysdale B-E, Grange PA, Cassels FJ, Wang L- X, Stamatos N, Farese A, Casey D, Powell J, Bhattacharjee AK, Kleinberg M, Goldblum SE. Recruitment of murine neutrophils in vivo through endogenous sialidase activity. J Biol Chem 2003;278:4112–4120.

48. Pshezhetsky AV, Hinek A. Where catabolism meets signalling: neuraminidase 1 as a modulator of cell receptors. Glycoconj J 2011;28:441–452.

49. Abdulkhalek S, Amith SR, Franchuk SL, Jayanth P, Guo M, Finlay T, Gilmour A, Guzzo C, Gee K, Beyaert R, Szewczuk MR. Neu1 sialidase and matrix metalloproteinase-9 cross-talk is essential for Toll-like receptor activation and cellular signaling. J Biol Chem 2011;286:36532– 36549.

50. Feng C, Stamatos NM, Dragan AI, Medvedev A, Whitford M, Zhang L, Song C, Rallabhandi P, Cole L, Nhu QM, Vogel SN, Geddes CD, Cross AS. Sialyl residues modulate LPS-mediated signaling through the Toll-like receptor 4 complex. PLoS One 2012;7:e32359.

51. Blander JM, Medzhitov R. Regulation of phagosome maturation by signals from toll-like receptors. Science 2004;304:1014–1018.

52. Doyle SE, O’Connell RM, Miranda GA, Vaidya SA, Chow EK, Liu PT, Suzuki S, Suzuki N, Modlin RL, Yeh W-C, Lane TF, Cheng G. Toll-like Receptors Induce a Phagocytic Gene Program through p38. Journal of Experimental Medicine 2004;

53. El-Benna J, Dang PM-C, Gougerot-Pocidalo M-A. Priming of the neutrophil NADPH oxidase activation: role of p47phox phosphorylation and NOX2 mobilization to the plasma membrane. Semin Immunopathol 2008;30:279–289.

54. Brinkmann V. Neutrophil Extracellular Traps Kill Bacteria. Science 2004;

55. Böhmer RH, Trinkle LS, Staneck JL. Dose effects of LPS on neutrophils-in a whole blood flow cytometric assay of phagocytosis and oxidative burst. Cytometry 1992;

56. Rifat S, Kang TJ, Mann D, Zhang L, Puche AC, Stamatos NM, Goldblum SE, Brossmer R, Cross AS. Expression of sialyltransferase activity on intact human neutrophils. J Leukoc Biol 2008;84:1075–1081.

57. Feng C, Zhang L, Almulki L, Faez S, Whitford M, Hafezi-Moghadam A, Cross AS. Endogenous PMN sialidase activity exposes activation epitope on CD11b/CD18 which enhances its binding interaction with ICAM-1. J Leukoc Biol 2011;90:313–321.

58. Sakarya S, Rifat S, Zhou J, Bannerman DD, Stamatos NM, Cross AS, Goldblum SE. Mobilization of neutrophil sialidase activity desialylates the pulmonary vascular endothelial surface and increases resting neutrophil adhesion to and migration across the endothelium. Glycobiology 2004;14:481–494.

59. Petri B, Phillipson M, Kubes P. The physiology of leukocyte recruitment: an in vivo perspective. J Immunol 2008;180:6439–6446.

60. Worthen GS, Schwab B 3rd, Elson EL, Downey GP. Mechanics of stimulated neutrophils: cell stiffening induces retention in capillaries. Science 1989;245:183–186.

61. Sheu C-C, Gong MN, Zhai R, Chen F, Bajwa EK, Clardy PF, Gallagher DC, Thompson BT, Christiani DC. Clinical characteristics and outcomes of sepsis-related vs non-sepsis-related ARDS. Chest 2010;138:559–567.

62. Wang T, Du Z, Zhu F, Cao Z, An Y, Gao Y, Jiang B. Comorbidities and multi-organ injuries in the treatment of COVID-19. Lancet 2020;

63. Wu Z, McGoogan JM. Characteristics of and Important Lessons From the Coronavirus Disease 2019 (COVID-19) Outbreak in China: Summary of a Report of 72 314 Cases From the Chinese Center for Disease Control and Prevention. JAMA 2020;323:1239–1242.

64. Hottz ED, Azevedo-Quintanilha IG, Palhinha L, Teixeira L, Barreto EA, Pão CRR, Righy C, Franco S, Souza TML, Kurtz P, Bozza FA, Bozza PT. Platelet activation and platelet-monocyte aggregate formation trigger tissue factor expression in patients with severe COVID-19. Blood 2020;136:1330–1341.

65. Wang J, Jiang M, Chen X, Montaner LJ. Cytokine storm and leukocyte changes in mild versus severe SARS-CoV-2 infection: Review of 3939 COVID-19 patients in China and emerging pathogenesis and therapy concepts. J Leukoc Biol 2020;108:17–41.

66. Vogl T, Stratis A, Wixler V, Völler T, Thurainayagam S, Jorch SK, Zenker S, Dreiling A, Chakraborty D, Fröhling M, Paruzel P, Wehmeyer C, Hermann S, Papantonopoulou O, Geyer C, Loser K, Schäfers M, Ludwig S, Stoll M, Leanderson T, Schultze JL, König S, Pap T, Roth J. Autoinhibitory regulation of S100A8/S100A9 alarmin activity locally restricts sterile inflammation. J Clin Invest 2018;128:1852–1866.

67. Liu J, Zhang S, Wu Z, Shang Y, Dong X, Li G, Zhang L, Chen Y, Ye X, Du H, Liu Y, Wang T, Huang S, Chen L, Wen Z, Qu J, Chen D. Clinical outcomes of COVID-19 in Wuhan, China: a large cohort study. Ann Intensive Care 2020;10:99.

