## Supplementary material for "Neuraminidase inhibitors rewire neutrophil function *in vivo* in murine sepsis and *ex vivo* in COVID-19": Supplem methods and refererences

**SUPPLEMENTARY METHODS**

**Human blood samples**

Blood was collected from healthy donors (25 - 45 yr old, n=3-12) in endotoxin-free tubes with K_3_EDTA (Labor Import, Brasil). All participants gave their written informed consent for blood collection after being informed on procedures. The research protocol followed the World Medical Association Declaration of Helsinki and was approved by the Institutional Review Board of the Federal University of Santa Catarina (CAAE #82815718.2.0000.0121). Blood samples were also collected from severe COVID-19 (n=6) or convalescent COVID-19 (n=8) patients (25 to 89 yr old) admitted in the Intensive Care Unit (ICU) or NUPAIVA (Research Center on Asthma and Airway Inflammation) at the UFSC University Hospital from August to October 2020. Severe COVID-19 patients (n=8) (49 to 93 yr old) in the Pneumology Department of Cochin Hospital, Assistance Publique Hôpitaux de Paris, France (February to April 2021) were also included. Blood samples from sex-matched healthy donors were used as controls. All patients or a close family member gave consent for participation in the study, which was approved by the Institutional Review Board of the UFSC (CAAE #36944620.5.1001.0121) and the Comité de Protection des Personnes Nord Ouest IV (ID-RCB 2020-A02700-39). Supplementary Table 1 summarizes patients clinical and laboratory records. These samples were used to analyze neutrophil activation, surface sialic acid as well as the effect of plasma under these parameters and ROS production.

**Evaluation of neutrophil activation, phagocytosis, killing, ROS, and NETs release**

Whole blood containing 1 x 10^6^ leukocytes were incubated (37 ^o^C, 5% CO_2_) in the presence or absence of Oseltamivir (100 µM, Sigma-Aldrich, San Luis, MO, USA), Zanamivir (30 µM, Sigma-Aldrich), LPS (1 µg/mL, *E. coli* 0127:b8, Sigma-Aldrich), LPS plus Oseltamivir or LPS plus Zanamivir for 90 min. Concentrations of Oseltamivir (100 µM) and Zanamivir (30 µM) showed in the experiments here were chosen by the best concentration-effect curves (10-100 µM Oseltavimir and 1-30 µM Zanamivir) with minimum effect on cell apoptosis/necrosis (95% of cell viability) (data not shown). Since plasma is a rich source of glycoconjugates, total leukocytes were used instead of whole blood to evaluate the effect of isolated neuraminidase from *Clostridium perfringens* (CpNEU) on neutrophils. Red blood cells (RBCs) were lysed by lysis buffer (0.15 M NH_4_Cl; 0.1 mM EDTA; 12 mM Na_2_HCO_3_) for 7 min, RT, followed by centrifugation (270 x g; 22°C; 7 min). Total leukocytes (1 x 10^6^ cells) were incubated (37 ^o^C, 5% CO_2_) in the presence or absence of CpNEU (10 mU, Sigma-Aldrich), CpNEU plus Oseltamivir (100 µM) or CpNEU plus Zanamivir (30 µM) for 60 min. Next, the following assays were performed. *Analysis of neutrophil activation*. Leukocytes were then washed and resuspended in flow cytometry (FACS) buffer (2 mM EDTA/PBS). The mix of antibodies against CD66b (G10F5; BioLegend, San Diego, CA, USA), CD62L (DREG-56; BioLegend), CD16 (3G8; BioLegend), isotypes or *Maackia amurensis* Lectin II biotinylated (MAL-II, Vector Labs, San Diego, CA, USA) coupled to Streptavidin (Biolegend),  *Sambucus Nigra* (Elderberry Bark) Lectin (SNA, Thermofisher), peanut agglutinin from *Arachis hypogaea* (PNA, Thermofisher)  plus human BD Fc Block^TM^ (BD Pharmingen^TM^) and Fixable Viability Stain (FVS, BD Horizon^TM^, San Jose, CA, USA) were added to leukocytes for 30 min at 4 ^o^C. Cells were washed, resuspended in FACS buffer, acquired in a FACSVerse cytometer and analyzed using FlowJo software (FlowJo LLC). Approximately 100.000 gated events were acquired in each analysis (Gate strategy shown in Supplementary Fig. 1). *Phagocytosis assays*. After RBCs lysis, 1 x 10^6^ leukocytes were incubated at 37 ^o^C (5% CO_2_) or at 4 °C (control) with 100 µg/mL pHrodo™ Red *E. coli* BioParticles® (Thermo Fisher, Waltham, MA, USA) for 60 min and the MFI of neutrophils (FVS^-^/CD66b^+^ cells) with ingested bioparticle was analyzed by FACS. Total leukocytes were also incubated with 1 x 10^6^ CFU of live *E. coli* (ATCC 25922) for 90 min (37 ^o^C, 5% CO_2_). Next, the cells were washed twice (2 mM EDTA/PBS), fixed (FACS buffer/PFA 2%) and the percentage of neutrophils with bacteria or the percentage of neutrophils with ≥3 bacteria was analyzed by light microscopy using Differential Quick Stain Kit (Laborclin, Brazil). *Bacterial killing*. Total leukocytes (1 x 10^6^) were incubated (37 ^o^C, 5% CO_2_) with 1 x 10^6^ CFU of live *E. coli* for 180 min. The samples were centrifuged (270 g, 7 min, 4 ^o^C) and 10 µL of supernatant were diluted until 10^6^ and spread onto agar brain-heart infusion (BHI, Kasvi, Brazil) to quantify the viable extracellular bacteria. The pellets were washed twice with PBS/2 mM EDTA (270 g, 7 min, 4 ^o^C), the leukocytes were lysed with 2% Triton-X, washed (PBS, 2000 g, 15 min, 4 ^o^C), resuspended in PBS and 10 µL of samples were diluted until 10^6^ and spread onto agar BHI. Plates were incubated overnight at 37 ^o^C and viable bacteria were expressed as mean ± SEM of CFU/mL. *ROS assay*. After RBCs lysis, 1 x 10^6^ leukocytes were incubated at 37 ^o^C (5% CO_2_) with 10 µM of cell-permeant 2',7'-dichlorodihydrofluorescein diacetate (CM-H2DCFDA, ThermoFisher) for 5 min. Next, cells were stimulated or not with phorbol 12-myristate 13-acetate (PMA) for 10 min, fixed, washed twice with PBS/2 mM EDTA (270 g, 7 min, 4 ^o^C) and analyzed by FACS. *NETs assay*. NETs quantification was performed as previously described (1) on the supernatant of isolated neutrophils. Briefly, an anti-MPO antibody bound to a 96-well flat-bottom plate captured the enzyme MPO (5 μg/ml; Abcam), and the amount of DNA bound to the enzyme was quantified using the Quant- iT™ PicoGreen® kit (Invitrogen, Carlsbad, CA, USA) according to the manufacturer's instructions. Fluorescence intensity (Ex 488 nm/Em 525 nm) was quantified in a FlexStation 3 Microplate Reader (Molecular Devices, San Jose, CA, USA). *Neutrophil isolation*. Human circulating neutrophils were isolated by Percoll density gradients (2). Briefly, four different gradients, 72%, 65%, 54%, and 45%, were used to isolate human circulating neutrophils. After centrifugation at 600 g for 30 min at 4 °C, the cell layer at the 72% gradient interface was collected as the neutrophil fraction. The erythrocytes were removed by lysis, and cell pellets were resuspended in RPMI 1640. Isolated neutrophils (1 x 10^6^/well) were treated with Oseltamivir, Zanamivir or medium 1 h before the stimulus with PMA (50 nM) or LPS (10 µg/mL). After 4 hr of stimuli (37 ^o^C, 5% CO_2_), the supernatant was collected to measure the levels of NETs.

**Neuraminidase kinetics assay**

After RBCs lysis, 0.5 x 10^6^ leukocytes were resuspended in HBSS and added to 96-well flat-bottom dark plate (SPL Life Sciences, South Korea) on ice. Then, 4- Methylumbelliferyl-N-acetyl-⍺-D-Neuramic Acid (4-MU-NANA, Sigma-Aldrich) substrate (0.025 mM) was added followed by medium, or LPS (1 µg/mL), LPS plus Oseltamivir (100 µM), LPS plus Zanamivir (30 µM). CpNEU (10 mU), CpNEU plus Oseltamivir or CpNEU plus Zanamivir were used as positive controls of the assay. The volume was completed to 200 μL with HBSS, followed by reading on the Spectramax® Paradigm® instrument starting 3 min after and every 5 min for 55 min at 37 ºC. Sialidase activity was also assessed in heat-inactivated or fresh plasma from severe COVID-19 patients in the presence or absence of Oseltamivir (100 µM) or Zanamivir (30 µM) using Tecan Infinite 200 multi-reader. The fluorescent substrate 4-MU-NANA formation was detected at ex 350 nm/em 450 nm.

**Immunofluorescence**

Isolated neutrophils (0.3 x 10^6^/slide) were fixed with 2% paraformaldehyde for 30 min and blocked with 5% BSA in PBS. Cells were then incubated for 1 hour at 37 °C with *Maackia amurensis* Lectin II biotinylated (15 µg/mL) diluted in 1% BSA/PBS. After washing with PBS, cells were incubated with Streptavidin Alexa Fluor 555 conjugate (1:1000; s32355 Life) for 30 min at 37 °C. Slides were prepared and nuclei stained with Mounting Medium containing DAPI (Sigma Aldrich). Stained cells were examined with a Widefield Zeiss fluorescence microscope, and the images were imported into ImageJ software for analysis.

**scRNA-seq Analysis**

Publicly available scRNA-seq data profiling nasopharyngeal or pooled nasopharyngeal/pharyngeal swabs (NSs), bronchial protected specimen brushes (PSBs) and bronchial lavages (BLs) were downloaded from https://figshare.com/articles/COVID-19_severity_correlates_with_airway_epithelium-immune_cell_interactions_identified_by_single-cell_analysis/12436517. Raw counts data, normalized data and UMAP coordinates generated by the authors of the original publication were imported into CellRouter for downstream analysis (3).

**Neutrophil responses with plasma from COVID-19 patients**

Whole blood samples from sex-matched healthy donors (n = 7) were incubated for 2 h (37 °C, 5% CO_2_) with 7% of fresh plasma from healthy donors, severe or convalescent COVID-19 patients or heat-inactivated plasma (56 °C, 30 min) from severe COVID-19 patients in the presence or absence of Oseltamivir (100 µM) or Zanamivir (30 µM). Surface levels of *α*2-3-Sia and CD66b and ROS production were assessed on neutrophils by FACS.

**Mice**

The care and treatment of the animals were based on the Guide for the Care and Use of Laboratory Animals (4) and all procedures followed the ARRIVE guidelines and the international principles for laboratory animal studies (5). C57BL/6 (Jackson Laboratory, Bar Harbor, ME, USA) mice (8–10 weeks old) and Swiss mice (10–12 weeks old) were housed in cages at 21 ± 2°C with free access to water and food at the Animal Facility of the Department of Microbiology, Immunology, and Parasitology and Department of Pharmacology from UFSC, respectively. A total of 228 mice were used in this study. Protocols were approved by the Animal Use Ethics Committee of UFSC (CEUA #8278290818).

***E. coli-, Klebsiella pneumoniae-* and CLP-induced sepsis**

*E. coli* (ATCC 25922, Manassas, VA, USA) or *K. pneumoniae* (ATCC 700603) were used for peritonitis- or pneumonia-induced severe sepsis in mice, as previously described (6, 7). Naive mice were intraperitoneal (IP) challenged with 100 µL of 1 x 10^7^ CFU of the *E. coli* suspension. A group of *E. coli*-septic mice was randomly pretreated (2 hr before infection) and post treated by *per oral* (PO, 12/12 hr) via with saline or Oseltamivir phosphate (10 mg/kg, Eurofarma, Brazil) for 4 days to survival analysis. Another group was pretreated 2 hr before infection with a single dose of Oseltamivir phosphate (10 mg/kg, PO) and the pathophysiological response was analyzed at 4 and 6 hr after infection. *E. coli*-septic mice were also randomly posttreated (6 hr after infection, 12/12 hr) with saline or Oseltamivir phosphate (PO, 10 mg/kg) for 4 days to survival analysis. For pneumonia-induced sepsis, mice were anesthetized with isoflurane (3–5 vol%) and placed in supine position. A small incision was made in the neck where the trachea could be localized and a *K. pneumoniae* suspension (4 x 10^8^ CFU/50 µL of PBS) was injected into the trachea with a sterile 30- gauge needle. Skin was sutured and animals were left for recovery in a warm cage. After 6 hr of infection and then 12/12 hr mice were treated with Oseltamivir phosphate (PO, 10 mg/kg) for survival analysis. In another set of experiments, pneumonia was induced and mice were treated 6 hr after infection with a single dose of Oseltamivir phosphate (10 mg/kg, PO) for material collection and analysis of pathophysiological response 24 hr after infection.

Cecal ligation and puncture (CLP)-induced sepsis were performed as previously described (8). Mice were anesthetized with xylazine (2 mg/kg, IP, Syntec, Brazil) followed by isoflurane (3–5 vol%, BioChimico, Brazil), a 1 cm midline incision was made in the anterior abdomen, and the cecum was exposed and ligated below the ileocecal junction. The cecum was punctured twice with an 18-gauge needle and gently squeezed to allow its contents to be released through the punctures. Sham-operated (Sham) animals underwent identical laparotomy but without cecal ligation and puncture. The cecum was repositioned in the abdomen, and the peritoneal wall was closed. All animals received 1 mL of 0.9% saline subcutaneous (SC) and 100 µL of tramadol (5 mg/kg, SC, Vitalis, Brazil) immediately after CLP. CLP-septic mice were randomly treated (starting 6 h after infection, PO) with 100 µL of saline or Oseltamivir phosphate (10 mg/kg, 12/12 hr) for 36 hr. In another set of experiments, CLP mice were randomly IP treated (6 hr after infection, 12/12 hr) during 4 days with 100 µL metronidazole (15 mg/kg, Isofarma, Brazil) plus ceftriaxone (40 mg/kg, Eurofarma, Brazil) and Oseltamivir phosphate (10 mg/kg) or saline by PO to survival analysis or treated for 36 hr to analyze the pathophysiological response at 48 hr after CLP.

**Neutrophil migration**

The animals were euthanized in a CO_2_ chamber, the bronchoalveolar lavage (BAL) and peritoneal lavage (PL) were performed and the number of neutrophils was determined at 4 and 6 hr after *E. coli*, 24 hr after *K. pneumoniae* infection or 48 hr after CLP surgery, as described^28^. Next, mice were perfused with PBS/EDTA (1 mM) and the lungs were harvested. Lungs were passed through 40-µm nylon cell strainers and single-cell suspensions were centrifuged in 35% Percoll® solution (315 mOsm/kg, Sigma-Aldrich) for 15 min at 700 g to enrich leukocytes populations. Pelleted cells were then collected, and erythrocytes were lysed. Single-cell suspensions from individual mice were determined using a cell counter (Coulter ACT, Beckman Coulter, Brea, CA, USA) or with a haemocytometer. Differential counts were also determined on Cytospin smears stained using Differential Quick Stain Kit (Laborclin, Brazil). Blood samples were collected by heart puncture and tubes containing heparin for further analysis. Neutrophils from LP or BAL were also stained with anti-Ly-6G/Ly-6C (GR-1, RB6-8C5; BioLegend) and MAL-II to be further analyzed by FACS, as previously described. Analysis was carried out in SSC^high^/GR-1^high^ cells.

**Bacterial counts**

The bacterial counts were determined as previously described (8). Briefly, the BAL, PL or blood were harvested and 10 µL of samples were plated on Muller-Hinton agar dishes (Difco Laboratories, Waltham, MA, USA) and incubated for 24 hr at 37 °C. PL or BAL samples were diluted until 10^6^.

**ELISA**

TNF (R&D Systems, Minneapolis, MN, USA) and IL-17 (XpressBio Life Sciences Products, Frederick, MD, USA) levels in plasma, PL or BAL were determined by ELISA kits according to the manufacturer’s instructions.

**Tissue injury biochemical markers**

Aspartate aminotransferase (AST), alanine aminotransferase (ALT), alkaline phosphatase (ALP) activities, and the levels of total bilirubin were determined in plasma samples by commercial kits (Labtest Diagnóstica, Brazil). The procedures were carried out according to the manufacturer's instructions.

**Statistical analysis**

The data are reported as the mean or median ± SEM of the values obtained from two to seven independent experiments. Each experiment using human samples was performed using three to five samples from healthy donors or one to three samples from severe or convalescent COVID-19 patients. We used five mice per experimental group except for survival analyses in which twelve to twenty mice were used. The mean or median values for the different groups were compared by analysis of variance (ANOVA) followed by Dunnett and/or Tukey post-tests. Bacterial counts were analyzed by the Mann–Whitney *U-*test or unpaired *t*-test using a parametric test with Welch’s correction. Survival curves were plotted using the Kaplan–Meier method and then compared using the log-rank method and Gehan-Wilcoxon test. Data was analyzed using GraphPad Prism version 8.00 for Mac (GraphPad Software, USA). A *P* < 0.05 was considered significant.

**Supplementary Table 1. Demographical, clinical and biological characteristics of patients (n=22)**

| ***Clinical characteristics at inclusion*** | |
| --- | --- |
| Gender (Male/Female) | 14/8 |
| Age (Years, Median (IQR)) | 51.7 (25 - 84) |
| Obesity (n, (%)) | 5 (22.7%) |
| Diabetes (n, (%)) | 8 (36.4%) |
| Cancer (n, (%)) | 4 (18.2%) |
| Immunodepression (n, (%)) | 2 (9.1%) |
| Hospitalization duration before inclusion (Days, Median (IQR)) | 3.5 (2.25 – 4.75) |
| Duration of symptoms before inclusion (Days, Median (IQR)) | 11 (10.25 – 13.25) |
| Dexamethasone treatment duration (Days, Median (IQR)) | 3 (2.25 – 4) |
| ***Oxygen dependance at inclusion*** | |
| Oxygen flow <3L/mn | 1 (4.5%) |
| Oxygen flow >3L/mn | 2 (9.1%) |
| Nasal high flow oxygen (n, (%)) | 5 (22.7%) |
| ***Overall severity during complete hospitalization*** | |
| Maximal oxygen flow during hospitalization <3L/mn | 0 (0%) |
| Maximal oxygen flow during hospitalization >3L/mn | 2 (9.1%) |
| Need for nasal high flow oxygen during hospitalization (n, (%)) | 6 (27.3%) |
| Out-hospital discharge | 6 (27.3%) |
| Transfer in ICU for OT intubation | 1 (4.5%) |
| Deceased | 1 (4.5%) |
| Complete hospitalization duration (Days, Median (IQR)) | 13 (8.25 – 27.25) |
| ***Laboratory data at inclusion*** | |
| Hemoglobin (g/dL, Median (IQR)) | 13.6 (11.38 – 15.7) |
| Hematocrit (%, Median (IQR)) | 40.8 (33.4 – 40.75) |
| Leukocytes (/mm^3^, Median (IQR)) | 8102 (4070 – 10660) |
| Neutrophils (/mm^3^, Median (IQR)) | 6852 (3355 – 10098) |
| Lymphocytes (/mm^3^, Median (IQR)) | 1452 (208 – 3728) |
| Monocytes (/mm^3^, Median (IQR)) | 408 (168 – 961) |
| Platelets (10^3^/mm^3^, Median (IQR)) | 233 (117 – 436) |
| Activated fibrinogen (g/L, Median (IQR)) | 5.47 (5.125 – 6.92) |
| ELISA D-dimers (ng/mL, Median (IQR)) | 1022 (559 – 3123) |
| CRP (mg/L, Median (IQR)) | 95.3 (22 – 214) |

IQR: interquartile range

**REFERENCES**

1.	Czaikoski PG, Mota JMSC, Nascimento DC, Sônego F, Castanheira FV e. S, Melo PH, Scortegagna GT, Silva RL, Barroso-Sousa R, Souto FO, Pazin-Filho A, Figueiredo F, Alves-Filho JC, Cunha FQ. Neutrophil Extracellular Traps Induce Organ Damage during Experimental and Clinical Sepsis. *PLoS One* 2016;11:e0148142.

2.	Mestriner FLAC, Spiller F, Laure HJ, Souto FO, Tavares-Murta BM, Rosa JC, Basile-Filho A, Ferreira SH, Greene LJ, Cunha FQ. Acute-phase protein alpha-1-acid glycoprotein mediates neutrophil migration failure in sepsis by a nitric oxide-dependent mechanism. *Proc Natl Acad Sci U S A* 2007;104:19595–19600.

3.	Lummertz da Rocha E, Rowe RG, Lundin V, Malleshaiah M, Jha DK, Rambo CR, Li H, North TE, Collins JJ, Daley GQ. Reconstruction of complex single-cell trajectories using CellRouter. *Nat Commun* 2018;9:892.

4.	National Research Council, Division on Earth and Life Studies, Institute for Laboratory Animal Research, Committee for the Update of the Guide for the Care and Use of Laboratory Animals. *Guide for the Care and Use of Laboratory Animals: Eighth Edition*. National Academies Press; 2011.

5.	Kilkenny C, Browne WJ, Cuthill IC, Emerson M, Altman DG. Improving bioscience research reporting: the ARRIVE guidelines for reporting animal research. *Osteoarthritis Cartilage* 2012;20:256–260.

6.	Czaikoski PG, Nascimento DC, Sônego F, de Freitas A, Turato WM, de Carvalho MA, Santos RS, de Oliveira GP, dos Santos Samary C, Tefe-Silva C, Alves-Filho JC, Ferreira SH, Rossi MA, Rocco PRM, Spiller F, Cunha FQ. Heme oxygenase inhibition enhances neutrophil migration into the bronchoalveolar spaces and improves the outcome of murine pneumonia-induced sepsis. *Shock* 2013;39:389–396.

7.	Yang WH, Heithoff DM, Aziz PV, Haslund-Gourley B, Westman JS, Narisawa S, Pinkerton AB, Millán JL, Nizet V, Mahan MJ, Marth JD. Accelerated Aging and Clearance of Host Anti-inflammatory Enzymes by Discrete Pathogens Fuels Sepsis. *Cell Host Microbe* 2018;24:500–513.e5.

8.	Spiller F, Orrico MIL, Nascimento DC, Czaikoski PG, Souto FO, Alves-Filho JC, Freitas A, Carlos D, Montenegro MF, Neto AF, Ferreira SH, Rossi MA, Hothersall JS, Assreuy J, Cunha FQ. Hydrogen sulfide improves neutrophil migration and survival in sepsis via K+ATP channel activation. *Am J Respir Crit Care Med* 2010;182:360–368.
